# Parallel channels for motion feature extraction in the pretectum and tectum of larval zebrafish

**DOI:** 10.1101/748913

**Authors:** Kun Wang, Julian Hinz, Yue Zhang, Tod R. Thiele, Aristides B Arrenberg

**Author notes:** These authors contributed equally (Kun Wang, Julian Hinz). Correspondence should be addressed to A.B.A.

## Abstract

Non-cortical visual areas in vertebrate brains extract different stimulus features, such as motion, object size and location, to support behavioural tasks. The optic tectum and pretectum, two primary visual areas, are thought to fulfil complementary biological functions in zebrafish to support prey capture and optomotor stabilisation behaviour. However, the adaptations of these brain areas to behaviourally relevant stimulus statistics are unknown. Here, we used calcium imaging to characterize the receptive fields of 1,926 motion-sensitive neurons in diencephalon and midbrain. We show that many caudal pretectal neurons have large receptive fields (RFs), whereas RFs of tectal neurons are smaller and mostly size-selective. RF centres of large-size RF neurons in the pretectum are predominantly located in the lower visual field, while tectal neurons sample the upper-nasal visual field more densely. This tectal visual field sampling matches the expected prey item locations, suggesting that the tectal magnification of the upper-nasal visual field might be an adaptation to hunting behaviour. Finally, we probed optomotor responsiveness and found that even relatively small stimuli drive optomotor swimming, if presented in the lower-temporal visual field, suggesting that the pretectum preferably samples information from this region on the ground to inform optomotor behaviour. Our characterization of the parallel processing channels for non-cortical motion feature extraction provides a basis for further investigation into the sensorimotor transformations of the zebrafish brain and its adaptations to habitat and lifestyle.

## Introduction

Visual receptive fields (RFs) are specific areas in the visual field where stimuli will alter the firing status of neurons (Spillmann, 2014). The ability of the visual system to extract useful information from the visual environment is directly related to the form, organization and diversity of neuronal receptive fields in the vertebrate visual system. Task-relevant visual features are processed in parallel channels in the brain (Nassi and Callaway, 2009), starting in the retina (Baden et al., 2016). The optic tectum and pretectum (superior colliculus and accessory optic system in mammals) receive direct input from direction-selective retinal ganglion cells (Giolli et al., 2006; Hunter et al., 2013; Robles et al., 2014) and encode visual stimuli moving in different directions (Wang et al., 2019). These evolutionarily ancient structures have developed to support navigation and orienting behaviour in zebrafish – which do not have a visual cortex – already soon after hatching in five-day-old larvae (Beck et al., 2004; Niell and Smith, 2005). While zebrafish are an important model organism for non-cortical vision research, the division of feature extraction tasks between tectum and pretectum is still largely unknown, and needs to be resolved. In particular their roles in feature extraction in relation to behavioural tasks are crucial for a mechanistic understanding of sensorimotor transformations in zebrafish.

In the accessory optic system (AOS)/pretectum of vertebrates, many neurons have large RFs with broad direction tuning curves (Britto et al., 1981; Grasse and Cynader, 1984; Masseck and Hoffmann, 2008; Simpson, 1984; Walley, 1967). Such large RFs should help the animal to distinguish wide-field optic flow from local motion and to estimate ego-motion. This computation is particularly important, because many vertebrates use the outcome to stabilize gaze and body position (Portugues and Engert, 2009; Rinner et al., 2005). In larval zebrafish, the optokinetic response (OKR) has been shown to rely on pretectal computation (Kubo et al., 2014), and the pretectum is involved in the optomotor response (OMR) as well (Naumann et al., 2016). In invertebrates, similar computations mediating OMR behaviour have been identified in the lobula plate, where horizontal system cells show large RFs with preferred directions matching the rotational optic flow around the yaw axis (Krapp et al., 2001). Optogenetic manipulation of these neurons evoked yaw optomotor behaviours in fixed and tethered flies (Busch et al., 2018; Haikala et al., 2013). In zebrafish, the pretectum contains further anatomical sub-divisions (Yanez et al., 2018), including structures likely involved in processing small visual stimuli during prey capture (Muto et al., 2017; Semmelhack et al., 2014) and a pretectal dopaminergic cluster providing input to the optic tectum (Tay et al., 2011). However, the RF properties of zebrafish pretectal neurons as well as their relation to optomotor behaviour are still largely elusive.

In contrast to the pretectum, in the zebrafish tectum RF sizes and locations have been measured before (Bergmann et al., 2018; Niell and Smith, 2005; Preuss et al., 2014; Sajovic and Levinthal, 1982; Zhang et al., 2011). Tectal neurons have relatively small RFs, conforming to the idea that tectal neurons detect small-size moving objects in the view field and are needed for hunting behaviour (Gahtan et al., 2005).

It is expected that each of the brain areas is adapted to the specific tasks and behaviours the animal executes in its environment. The behavioural relevance of a visual stimulus is likely influenced by the visual field location of the stimulus and will depend on the particular visually mediated behaviour and the probability of observing such stimulus locations under natural conditions. For example, in the brain of macaque monkeys, it has been shown that RF properties in the superior colliculus, the homolog of the optic tectum, widely differ in the upper and lower visual view field, which likely represents adaptations to near-space in the lower and far-space in the upper visual field (Hafed and Chen, 2016). During hunting behaviour, vertebrates typically keep the (small) prey stimuli in their nasal/frontal visual field and therefore a high visual acuity in the nasal/frontal view field is advantageous. Accordingly, both the primary visual cortex and the superior colliculus show a magnification of foveal visual field regions (Grujic et al., 2018; Schwartz, 1980), i.e. more neurons are dedicated to represent these foveal locations than more peripheral locations. Also in the zebrafish retina, a region of heightened photoreceptor density has been described (area centralis (Schmitt. and Dowling., 1999)), corresponding to upper-nasal visual field positions (Zimmermann et al., 2018). For visual stabilization behaviours, animals need to detect their ego-motion. In order to use brain resources efficiently, the reliable detection of optic flow directions is likely biased towards making use of the most informative visual field locations that occur in natural habitats and during behaviour. The optomotor response is driven effectively by whole-field motion but – to our knowledge – there are no previous reports on particular visual field regions being preferably sampled by the animal to initiate OMR. Given the different roles of the optic tectum and the pretectum in hunting and stabilization behaviour, respectively, it seems likely that these brain areas represent the visual field differently. It is unclear, however, whether the observed retinal anisotropies are relayed to primary visual areas in the zebrafish brain and whether magnifications of certain visual field locations exist in the tectum or pretectum of zebrafish. The characterization of such brain-area specific magnifications within the small vertebrate brain of larval zebrafish would advance our understanding of the efficient encoding of relevant information in the vertebrate brain and help to reveal the specific computations that brains have evolved to perform.

Here we characterise the RF properties of tectal and pretectal motion-sensitive neurons using *in vivo* 2-photon calcium imaging of GCaMP5G transgenic animals and investigate their organisation in visual and anatomical space. In addition, we investigate how the identified RFs match to the visual locations which drive the OMR behaviour. Our results show the complementary roles of the optic tectum and pretectum to support behaviourally relevant motion feature extraction.

## Results

To estimate RF properties of pretectal and tectal neurons, we stimulated the right eye of immobilized larval zebrafish with a series of horizontally moving grating patterns of different sizes and locations (Figures 1A and S1, STAR Methods) and measured GCaMP5G calcium responses of neurons in the diencephalon and midbrain (Figure 1B). 1,926 motion-sensitive neurons that responded reliably during the three repetitions of the stimulus protocol were recorded in 10 animals. Neurons were divided into four functionally defined groups (Figures 1C-D, Supplementary Figure S2, see STAR method for classification), based on the size and shape of their receptive fields (RFs): (1) small-size RF, (2) medium-size RF, (3) large-size RF, and (4) bar-shaped RF. RF sizes ranged from very small RFs (30° x 13°) to whole field (168° azimuth x 80° elevation). In neurons with smaller RFs, we oftentimes observed suppressive effects for larger motion stimuli (Figure 1Di), showing that these neurons were small-size selective. Small-size RF neurons without signs of inhibition were frequently encountered as well: even though the excitatory RF density (STAR Methods) was localized to a small patch in the visual field, the neurons were also responsive to whole-field stimuli (Figure 1Dv). Furthermore, some of the small-size and medium-size RF neurons each responded to small moving stimuli in a range of different visual field positions, while not responding to larger moving stimuli covering the same visual field locations, i.e. their responses were small-size selective and position-invariant (Figure 1Dvi).

**Figure 1.**
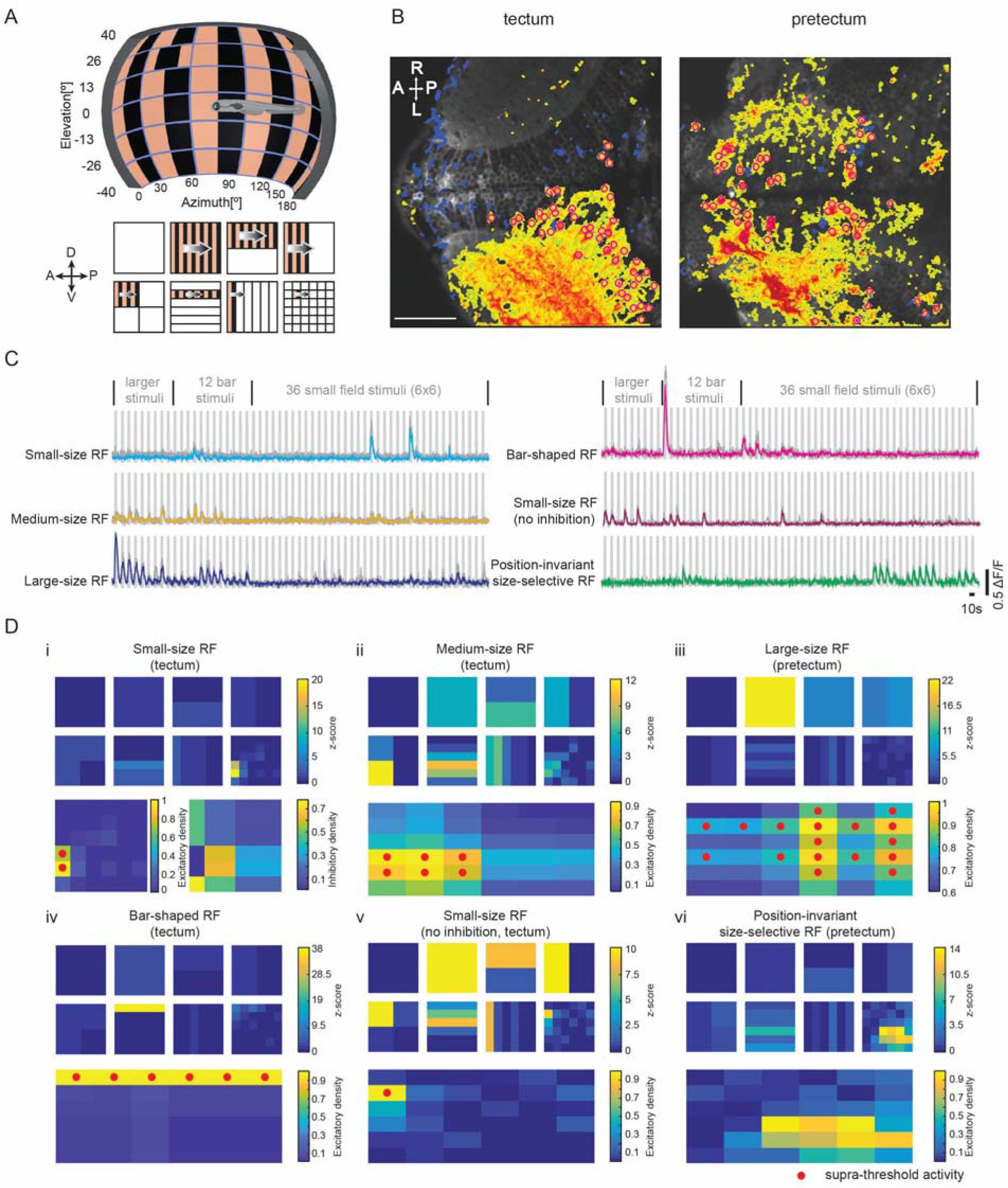
Different receptive field types were identified using horizontally moving gratings. **(A)** Stimuli of naso-temporally moving gratings covering view fields of variable sizes were presented to the animal’s right eye (n = 8 fish for naso-temporal and n = 2 fish for temporal-nasal motion). Motion stimuli consisted of whole-field (1×1, 180° x 80°, azimuth x elevation), half field (2 x 1/2 and 1/2 x 2), quarter field (2 x 2 x 1/4), bar (6 x 1/6 and 1/6 x 6) and small field (6 x 6 x 1/36, each 30° x 13°) stimuli. The non-stimulated regions are shown in white for illustration purposes but contained a stationary grating. A snapshot during the small field motion phase (depicted in the lower right) is shown on the display in the setup illustration (top). (**B**) The average fluorescence of example calcium imaging time series recorded from tectum and pretectum is shown in grey. Motion-sensitive image pixels are shown in false colour, with warm and cold colours corresponding to positive and negative motion phase correlation, respectively. Manually selected ROIs are labelled with magenta circles. Scale bar, 50 um; A, anterior; P, posterior; L, left; R, right. **(C)** Example ΔF/F calcium responses of neurons with different receptive field sizes or shapes are shown. For each neuron, the coloured trace corresponds to the median response across three repetitions (grey traces). The grey rectangular shades correspond to the 57 presented motion phases. **(D)** Receptive field maps for six example neurons corresponding to (C). Top: The eight squares correspond to the eight stimulus segments shown in (A) and each square corresponds to the stimulus arena surface (180° x 80°, azimuth x elevation). The calcium response is plotted as a z-score for each stimulus phase (for each ROI, the ΔF/F, subtracted by the average of the ΔF/F, divided by the standard deviation of the baseline ΔF/F). Bottom: By comparison of the activities evoked by spatially overlapping stimuli of different sizes, excitatory receptive field densities were calculated to measure the size of the receptive fields as the number of patches with supra-threshold activity (red dots). For cells with small-size excitatory receptive fields with maximal responses during the small-size stimulus phases (see STAR methods), an inhibitory receptive field density was calculated to judge the extent of small-size selectivity (cell i). The anatomical location of each neuron is indicated (tectum or pretectum).

### Pretectal RFs are larger than tectal RFs and they are less often size-selective

For each motion-sensitive neuron, we measured the location of the RF centre in the visual field and the anatomical position of the soma in the brain (see STAR Methods). We then investigated the topography of pretectal and tectal neurons as well as their sampling of the visual field. On average, we identified 205 ± 6 motion-sensitive neurons per tectum and 59 ± 13 neurons per pretectum (n = 10 fish, corresponding to 5 complete composite brains sampled in 10 µm steps).

Pretectal neurons have larger excitatory receptive fields as compared to tectal neurons (Figures 2A and 2B). Within the pretectum, 30% (88/295) of the motion-sensitive neurons had a large-size receptive field, compared to less than 2% (19/1,251) of the neurons within the tectum (Figure 2B). Most motion-sensitive tectal neurons (86%, Figure 2B) had excitatory receptive fields smaller than 1200 deg^2^ in area (3 of our small stimulus patches, each covering 30° x 13° in azimuth and elevation), which is consistent with previous reports from other groups (Bergmann et al., 2018; Sajovic and Levinthal, 1982). Furthermore, similar to the findings in a previous report (Preuss et al., 2014), we found that 68% (853/1,251) of motion-sensitive neurons in the tectum were size-selective, i.e. motion stimuli that were larger than the neuron’s excitatory RF evoked lower calcium responses (Figure 1Di and Figure 2B). In contrast, only 26% (77/295) of the motion-sensitive neurons in the pretectum were size-selective (Figure 2B).

**Figure 2.**
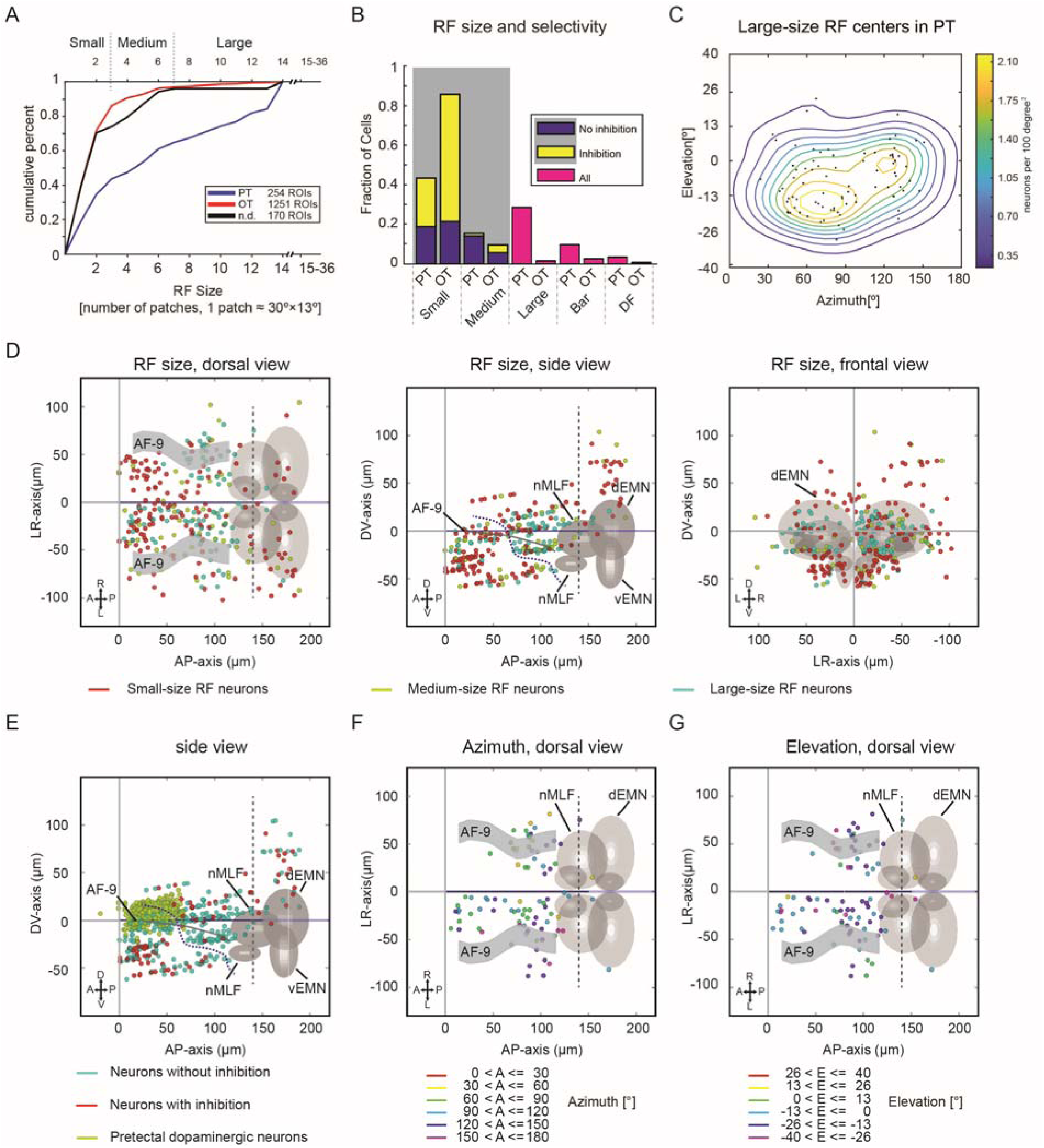
Pretectal receptive fields are large and biased towards the lower view field. **(A)** Analysis of receptive field size differences across pretectum and optic tectum. The cumulative distribution of receptive field sizes is shown for pretectum (blue), optic tectum (red) and neurons of undefined provenance (black). Note that the pretectum has a larger fraction of large-size RF cells compared to the tectum. **(B)** For each brain area (PT: pretectum; OT: optic tectum), the fractions of small-size, medium-size, large-size, bar-shaped, and double-field (DF) receptive fields are shown. For small- and medium-size RFs, the inhibitory surround was investigated and neurons with such inhibition are plotted in yellow. DF RF classification served as a control for stimulus reflections in our experiments (see Methods). **(C)** Locations and density contour plot of receptive field centres of large-size RF pretectal neurons in the contralateral and ipsilateral hemispheres (n = 10 fish, 5 composite brains). **(D)** Anatomical map of small-size (red), medium-size (yellow) and large-size RF (cyan) neurons in the pretectal region. (n = 10 fish, 5 composite pretectal regions, Figures S4B to S4D). Each coloured dot represents a single neuron. Coordinates are defined as distances relative to the posterior commissure in the diencephalon (anterior-posterior axis and dorso-ventral axis) and midline (left-right axis). **(E)** Anatomical map of three types of neurons in the pretectal region: neurons with (red) and without (cyan) signs of inhibition, and pretectal dopaminergic neurons (green). The dopaminergic neurons served as a landmark for the dorsal periventricular pretectal nucleus and were recorded in a separate experiment. **(F, G)** Topographic maps of large-size RF neurons in pretectal region. Each coloured dot represents a single neuron with its receptive field centre in the indicated azimuth **(F)** and elevation **(G)** range. For example, in panel (F), all receptive field centres of the neurons in red are located in the nasal-right side in front of the fish (0° to 30° azimuth). In panel (G), all receptive field centres of the neurons in green are located slightly above the equator of the view field (0° to 13° elevation) (n = 10 fish, 5 composite pretectal regions). dEMN, dorsal extraocular motor neurons; vEMN, ventral extraocular motor neurons; dEMN and vEMN, the trochlear and oculomotor nuclei; nMLF, nucleus of the medial longitudinal fasciculus; A, anterior; P, posterior; D, dorsal; V, ventral, L, left; R, right; AF, arborisation field. The abbreviations are applicable to all anatomical maps in this study. Neurons located ≥140 um caudal to the posterior commissure are located outside of the pretectal region and were excluded from pretectal analysis (dashed grey line).

### Large-size RFs in the caudal pretectum are biased to the lower visual field

The receptive field centres of 69% of the large-size RF pretectal neurons were located in the lower visual field, which represents a significant bias (Figure 2C and Supplementary Figure S1B, p = 0.0002, z-test for one proportion). These pretectal large-size RF neurons were located almost symmetrically in both hemispheres of the caudal pretectum, with some neurons in the rostral pretectum on the contralateral side (laterality index = −0.30, Figure 2D). The high number of ipsilateral neurons can – in part – be explained by reflections of the stimulus, which was revealed in an additional experiment in which the left eyes of the fish were blocked by a back foil (laterality index = −0.86; n = 6 fish, 6 pretecta; Figure S1C). In addition to the large-size RF neurons observed in the caudal pretectum, many neurons responsive to small moving stimuli were also identified in the pretectum (Figure 1Cvi, Figure 2D), particularly in the rostral region, which – to our knowledge – has not been described before.

Since neurons in the rostral pretectum segregated from those in the caudal pretectum through an anatomical gap containing only few motion-sensitive neurons (Figure 2D, side view), we defined a boundary based on this gap to separate these two anatomical clusters (dashed line in Figure 2D-E, Figure S4A and STAR Methods). The rostral pretectum contained a higher proportion of small-size RFs (57%, 85/148) than the caudal pretectum (29%, 43/147, p<0.001, z-test for two proportions), while the caudal pretectum contained a higher proportion of large-size RFs (43%, 63/147) than the rostral pretectum (14%, 21/148, p<0.001) (Figure 2D, S1C). The rostral pretectum furthermore contained a higher proportion of small-size selective neurons (Figure 2E, S4B-D), whose activity was suppressed for larger stimuli (rostral pretectum: 34% (51/148), caudal: 18% (26/147), n = 148 and 147, p<0.001, z test for two proportions, see STAR methods). Rostral pretectal neurons responsive to our smallest grating stimuli were also frequently (63% of the neurons, n = 6 fish) responsive to small horizontally moving dots of variable diameters (3 to 18 degrees diameter), as we tested in a separate experiment (data not shown).

It is well established that soma positions of zebrafish tectal neurons are topographically arranged within the tectum and that the receptive field centres cover almost the whole visual field at the population level (Attardi and Sperry, 1963; Bergmann et al., 2018; Niell and Smith, 2005; Romano et al., 2015). However, in the pretectum, we did not observe a clear topographic distribution of large-size RF neurons (Figures 2F-G and S1D-E).

### Tectal receptive fields dominate the upper-nasal view field

As expected from the topographic retino-tectal projection of retinal ganglion cells, strong rostro-caudal, dorso-ventral and medio-lateral topographical gradients of receptive field centres were found for small-size RF neurons in the optic tectum (Figures 3B, 3D, 3E and Supplementary Figure S3A). Responding tectal neurons were mainly located in the contralateral hemisphere relative to the stimulated eye (laterality index = −0.88, Figures 3D-E, see STAR Methods). More tectal neurons with small-size receptive fields corresponding to the upper-nasal visual field were found than corresponding to the lower-temporal visual field (420 vs 55 neurons, upper-nasal vs lower-temporal view field, z-score test for proportions: p<0.001, Figures 3A and 3C). When comparing the upper-nasal view field (26° elevation, 30° azimuth) to the lateral view field (0° elevation, 90° azimuth), the sampling difference corresponded to a 1.6 fold magnification factor (Figure S3B). In our experiments we identified about 4 tectal small-size RF neurons per 10°x10° in the upper-nasal field in each completely sampled fish (hypothetical sampling every 5 µm in the dorso-ventral direction, calculated based on our actually recorded 33.4 tectal motion-sensitive neurons per 60°x27° in each of 5 composite brains sampled every 10 µm, see Figures S3B-D).

**Figure 3.**
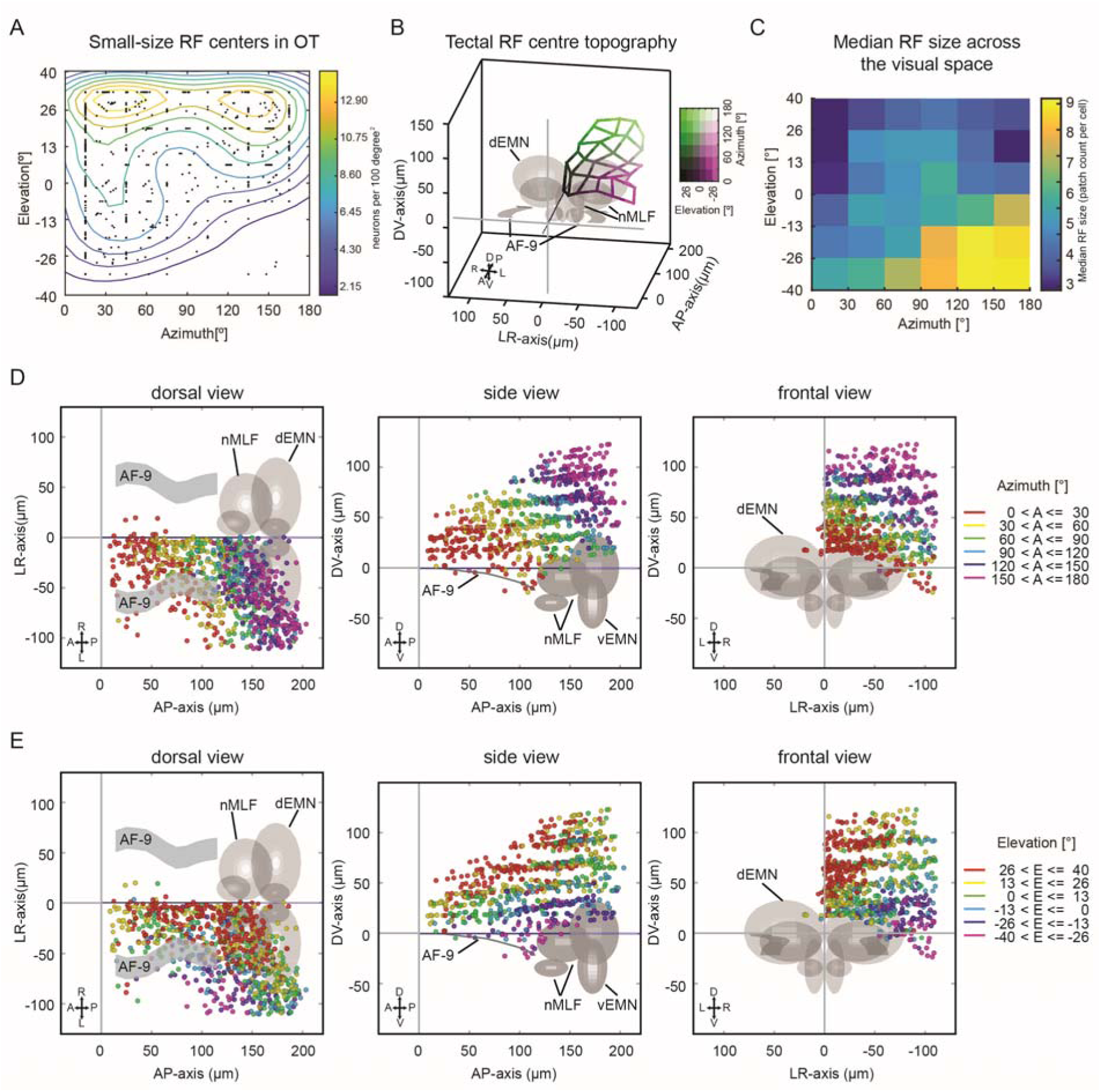
Tectal receptive fields are small and biased towards the upper nasal visual view field. **(A)** Visual field locations and density contour plot of receptive field centres of small-size RF tectal neurons (n = 10 fish, 5 composite tecta). **(B)** Anatomical map of the tectal receptive field centre topography. The average anatomical position of neurons with their receptive field centres in each of the 6 x 6 visual field bins was calculated and all 36 locations were connected by a grid to illustrate the mapping of visual space (colour legend in the upper right) corresponding to anatomical space in the tectum (Supplementary Figure S3A). The yaw angle (10° right) and the pitch angle (20° down) were adjusted to allow optimal view of the anatomical topography. **(C)** Median receptive field size across the visual space. For each patch, we calculated median receptive field sizes of all neurons (in tectum and pretectum) whose excitatory RFs covered the patch in question. The animals sample the lower-temporal visual field mainly with large-size RF neurons, while small-size neurons dominate in the upper nasal visual field. (**D, E**) Topographic maps of tectal small-size RF neurons for azimuth (D) and elevation (E). Each coloured dot represents a single neuron with its receptive field centre in the corresponding azimuth range in **(D)**. For example, all receptive field centres of the neurons in red are located between 0° azimuth (in front of the fish) and 30° azimuth on the nasal right side of the fish. In (**E**) each coloured dot represents a single neuron with its receptive field centre in the corresponding elevation range. For example, receptive field centres of the neurons in green are located slightly above the equator of the view filed (0° to 13° in elevation). n = 10 fish, 5 composite brains.

### Zebrafish OMR behaviour is driven best by motion in the lower-temporal visual field

While it is known that during hunting behaviour, prey stimuli are mostly located in nasal visual field locations in zebrafish (Bianco et al., 2011), the visual field regions that drive OMR and OKR behaviour have not been identified. OMR behaviour can be stably induced with whole-field forward motion projected from below or from the side in larval zebrafish (Severi et al., 2014; Thiele et al., 2014), and it had been assumed that large – or even – whole-field stimuli are necessary to drive stabilisation behaviours. Given the uneven distribution of large-size RFs in the upper and lower visual fields in the pretectum (Figure 2C), we wanted to test whether OMR is mainly driven by the forward motion located in the lower visual field, which would implicate the large-size RF neurons in mediating the OMR behaviour. We therefore recorded the tail motion of larvae while forward-moving gratings of different sizes were presented in different visual field locations. Care was taken to always stimulate animals in a binocularly symmetric fashion to drive forward OMR instead of OMR turning behaviour (Figures 4A-B and S5A-B). The angle between the anterior-posterior body axis and the tail tip was traced to detect single tail beats and swim bouts (Figures 4A, 4D and 4E). Bouts, consisting of series of tail undulations beating symmetrically to both sides (forward OMR), were induced in response to whole-field forward moving gratings (Supplementary Movie S1). In contrast, unsymmetrical unilateral turning swim beats to one side (turning OMR) were oftentimes evoked by whole-field rotating visual stimuli, although symmetrical beats (forward OMR) were also observed (Figure S5Ei). Whole-field and half-field visual stimuli covering the temporal or lower view field could evoke forward OMR robustly (Figure 4C).

**Figure 4.**
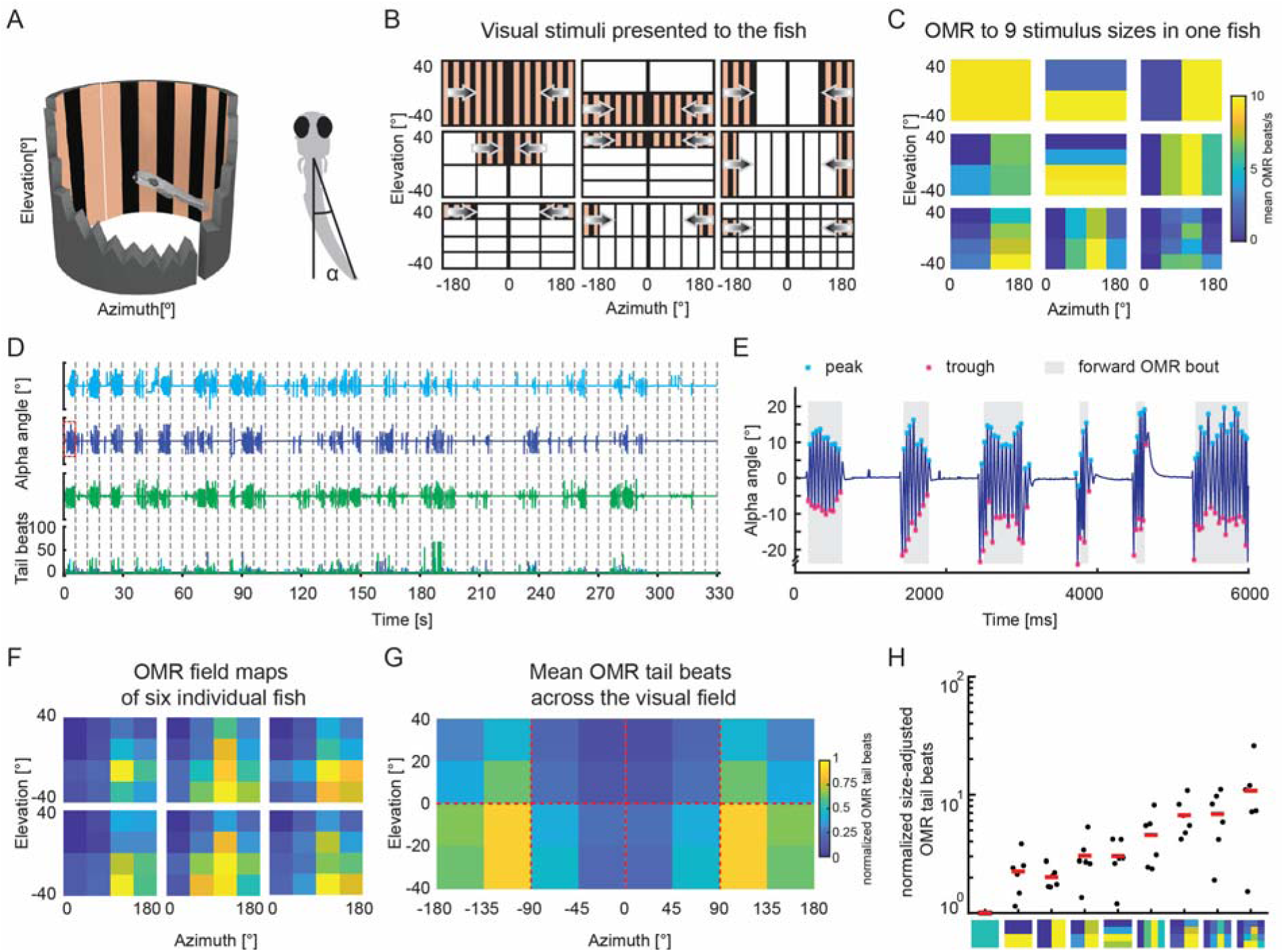
Zebrafish optomotor responses are driven by motion from below and in the rear of the animal. **(A)** Left: stimuli were presented binocularly to the animal with two half-cylindrical arenas while the fish behaviour was recorded. A snapshot of the visual stimulus during the whole field motion phase is shown in the setup illustration. Right: a schematic of a larval zebrafish performing OMR behaviour. The indicated angle between the anterior-posterior body axis and the tail tip was measured to judge OMR performance. **(B)** Motion stimuli of the OMR experiment consisted of whole-field (1×1, 180° azimuth x 80° elevation), half field (2 x 1/2, 1/2 x 2), quarter field (2 x 2 x 1/4), bar (4 x 1/4, 1/4 x 4) medium field (4 x 2 x 1/8 and 2 x 4 x 1/8) and small field (4 x 4 x 1/16, each 45° x 20°) stimuli (for each size group, only one motion patch is illustrated). Each stimulus was mirror-symmetric about the mid-sagittal plane and shown with two half cylindrical arenas from both sides. The control stimulus phases (not shown in panel (B)) consisted of two stationary whole-field phases, counter-clockwise and clockwise rotational moving gratings, or looming visual stimuli on either side, Supplementary Figure S5B). The regions in which no stimulus movement was present are shown as white areas for illustration purposes but contained a stationary grating. **(C)** Normalized number of tail undulations per second for one larva, induced by forward moving gratings of 9 different sizes (rectangles represent the stimulus hemi-fields from (B)). **(D)** The full behavioural session for one animal, showing induced OMR behaviour by stimuli of different sizes and locations as indicated in panel (B). The three stimulus repetitions are shown in different colours (cyan, blue and green). Individual stimulus phases are separated by the dashed grey lines and each row corresponds to 55 concatenated time periods in which motion stimuli were presented. The stimulus pauses in-between motion phases were cropped out and lasted 69 s each. Measured tail beat counts (bottom) during each trial were consistent across the three repetitions. **(E)** Tail movements induced by whole-field forward moving gratings (from trial no. 2 indicated by a red rectangle in panel (D)). The peaks and troughs of each swim bout within the 6-s recording are labelled with cyan and magenta asterisks. OMR swim beats are indicated by grey background shade. **(F)** Visual field heat maps of the forward-OMR tail-beat rate for six individual larval zebrafish (n = 6). The visual field density of OMR behaviour was quantified analogous to the excitatory RF densities in Figure 1.The density was normalised according to the size of the stimulation area in each fish individually (see STAR Methods) (**G**) Average heat map of the OMR beat density for stimulation across different visual field coordinates (n = 6 fish). (**H**) Maximum OMR beats for each of the 9 stimulus fields shown in (Figure 4B-C) after normalization according to stimulus size (see STAR Methods).

To our surprise, forward OMR swim beats could be induced by stimuli as small as 45° x 20° (azimuth x elevation, the smallest size in our protocol) in the lower-temporal view field of both eyes (Figure 4C). The OMR-evoking visual field locations were almost identical across all recorded animals (n = 6, Figure 4F). To test whether the optomotor responses evoked by small stimuli are stronger than what would be expected under the assumption that OMR drive was established by the sum of equal-sized motion inputs across the visual field, we normalized the evoked OMR tail-beat rate to the respective stimulus field size (analogous to the excitatory RF density estimation, see STAR Methods). The resulting visual field map of OMR drive (Figure 4G) shows the disproportionally large influence of moving stimuli in the lower-temporal view field. We then compared OMR drive across different stimulus sizes (1×1, 2×1, … 4×4), always taking into account the visual field positions/stimulus phases which drove OMR best. This analysis revealed that the smallest stimulus area evoked responses, which were on average ∼11 times stronger than expected by an equal integration of optic flow inputs across the visual field (Figures 4H). The visual field locations mediating strong OMR responses matched qualitatively with the measured RF centers of caudal pretectal large-size RF neurons (Figure S5D), suggesting that large-size RF neurons in the caudal pretectum drive OMR behaviour.

## Discussion

Our study reveals the functional segregation of visual motion processing in parallel channels each extracting different sets of motion features across the visual field. The optic tectum, which processes motion of small visual stimuli, has a bias for upper-nasal visual field locations. The pretectum processes wide-field optic flow mainly in the caudal pretectum using large RFs mainly sampling the lower visual field, and processes small-size motion stimuli mainly in rostral regions. Furthermore, we show that animals observe mainly the lower-temporal visual field for optomotor forward swimming.

These findings are in agreement with the need to process small visual stimuli, e.g. during prey capture (Bianco et al., 2011; Preuss et al., 2014), and the need to assess wide-field motion to inform stabilization behaviours (Kubo et al., 2014). The data supports a circuit model in which these two distinct tasks are processed independently via multiple channels in different brain areas.

### Caudal and rostral pretectal regions are biased towards the encoding of large-field optic flow and small stimuli, respectively

Large-size RF pretectal neurons are mainly found in the caudal pretectal region, while small-size RF pretectal neurons are biased towards more rostral anatomical locations. The large-size RFs of pretectal neurons preferably sample the lower half of the visual field (Figures 2C and S1B), which fits with previous reports from pretectal neurons in dogfish (Masseck and Hoffmann, 2008).

Due to our use of a half-cylindrical stimulus arena to present exclusively horizontally moving stimuli to the right eye, neurons with more complex, e.g. rotational, binocular, or vertical optic flow fields could not be described in this study (Kubo et al., 2014; Wang et al., 2019). These types of RF structures exist in visual neurons of other species (Karmeier et al., 2003; Krapp et al., 2001) and future studies are needed to identify them in zebrafish.

The adult zebrafish pretectum contains several nuclei distributed from the superficial to the periventricular regions, receives numerous retinal and tectal afferents, and projects to the optic tectum as well (Fernald and Shelton, 1985; Kastenhuber et al., 2010; Presson et al., 1985; Yanez et al., 2018). The correspondence of these nuclei to functionally defined larval brain regions is not fully resolved (Arrenberg and Driever, 2013; Kubo et al., 2014; Muto et al., 2017; Semmelhack et al., 2014). In our study, the pretectal dopaminergic neurons, which are evolutionarily conserved across most amniotes (Yamamoto and Vernier, 2011), were used as landmark to indicate the location of the periventricular pretectal nucleus (Filippi et al., 2014). The location of caudal pretectal neurons in this study corresponds to the anterior medial cluster (AMC), previously described by Kubo et al. (Kubo et al., 2014). Recorded extra-tectal neurons located more than 140 um caudal to the posterior commissure most likely belonged to the tegmentum and were therefore excluded from the pretectal analysis (see STAR Methods). The rostral pretectal neurons described in this study mainly cover a rostro-ventral pretectal or potentially dorsal thalamic brain region (Rupp et al., 1996; Yanez et al., 2018). Functional properties of two larval pretectal nuclei have previously been described in relation to prey capture (parvocellular and magnocellular superficial nuclei, Psp and Psm), however they are located more laterally than the bulk of our recorded motion-sensitive neurons in the rostral pretectum (Muto et al., 2017; Semmelhack et al., 2014). Morphologically, the identity, extent, and overlap of larval pretectal neuron populations, which give rise to each of the known adult pretectal nuclei, is not easily discernible (Arrenberg and Driever, 2013). Further anatomical studies are needed to link our functionally identified neurons to specific pretectal brain nuclei and connectivity in the larval brain.

While the anatomical and visual field locations as well as neuron numbers were consistent for large-size RF neurons in the pretectum across fish and experiments (Figure 2C-D and Figure S1B-C), the responses of position-invariant RF neurons (Figure 1Cvi) appeared to be more variable. Notably, the preferred visual field locations, exact pretectal anatomical locations, and the number of identified position-invariant neurons within the pretectum differed in the first and second experiment we performed (Figure S4E-F). Further work is needed to elucidate the specific anatomical distribution and response properties of position-invariant RF neurons.

### The tectum – poised for prey capture

Our analysis of a large number of tectal neurons extends previous reports on tectal physiology (Niell and Smith, 2005; Preuss et al., 2014; Sajovic and Levinthal, 1982; Zhang et al., 2011). Our finding that the tectum mostly comprises relatively small excitatory RFs, which oftentimes are small-size selective, is in agreement with a role of the tectum in prey capture (Bianco et al., 2011; Gahtan et al., 2005). It is noteworthy that our stimulus protocol only allowed measurement of RF sizes down to 30° horizontally and 13° vertically (corresponding to our smallest visual stimulus), which is much larger than the minimal RF size in a previous report (Preuss et al., 2014). Also, due to the limitation of 2-phtoton calcium imaging, we are not able to directly assess inhibition, but assess it only indirectly by comparing the reduction in responses evoked by larger stimuli.

The topographic arrangement of the zebrafish tectal neurons is well established both for the anatomical retino-tectal projection (Baier et al., 1996; Trowe et al., 1996) and the functional RF mapping (Bergmann et al., 2018; Niell and Smith, 2005). However, precise measurements of dedicated tectal anatomical volumes had not been performed previously and are needed to build faithful models of zebrafish vision. We find that the upper nasal visual field (134 µm^3^/deg^2^) is magnified in the tectum by a factor of 1.6 relative to lateral (82 µm^3^/deg^2^) visual field locations (Figure S3), which is a relatively mild magnification in comparison to foveal magnification in the primate superior colliculus and visual cortex (Cowey and Rolls, 1974; Grujic et al., 2018; Schwartz, 1980). Given that the larval prey capture behaviour depends on the optic tectum and the larvae respond to the paramecia located in front and extending 60° temporalwards of the fish (Bianco et al., 2011; Romano et al., 2015), a role of these tectal small-size RF neurons in prey capture seems likely. Since the density of small-size RF neurons is elevated in the upper nasal view field (Figure 3A and S3B), this would suggest that larval prey capture performance is best when the paramecia are located in front and slightly above the eyes, and possibly the mouth. In this study, we have only investigated the RF distributions at a single developmental stage. It is possible that the reported tectal magnification of the upper-nasal visual field is (in part) a result of the tectal developmental stage, since new, initially non-functional neurons are added in the dorso-medial and caudo-lateral tectum (Boulanger-Weill et al., 2017; Recher et al., 2013).

### The optomotor response is driven strongest by the lower-temporal visual field

While it had been known from previous reports that motion stimuli presented from the bottom or from the side are effective in triggering OMR behaviour (Orger et al., 2008; Severi et al., 2014; Thiele et al., 2014), it was unclear to which part of the visual field the animal pays attention. We show that for forward swimming OMR, the relevant region lies in the lower-temporal view field of the fish (Figures 4F and 4G). The distribution of OMR drive matches qualitatively with the location of large-size RFs of the pretectal neurons (Figure S5D), which is in agreement with the hypothesized role of the pretectum in mediating OMR behaviour. Particularly, moving stimuli, as small as 45°x20° (azimuth x elevation), were able to evoke robust OMR behaviour in our experiment, even in the presence of the surrounding whole-field stationary stimulus of our stimulus protocol. This finding is in contrast to the concept that the OMR is a whole-field-induced behaviour. Rather, it suggests that distinct parts of the visual field are sampled for body stabilization behaviours. This most likely reflects the ecological adaptions of zebrafish living in shallow waters (Arunachalam et al., 2013) and the need to sample the most relevant parts of the visual field for different tasks (Zimmermann et al., 2018), such as putative high contrast textures in the river bed.

Since RGCs project directly to optic tectum and pretectum (Robles et al., 2014), it seems likely that the small-size RFs without inhibition as well as the large-size RFs in the caudal pretectum are established by direct inputs from RGCs (Figure 5). Small-size-selective responses require inhibitory inputs, which could be calculated already within the retina or within the retino-recipient brain areas (Grama and Engert, 2012; Ramdya and Engert, 2008).

**Figure 5.**
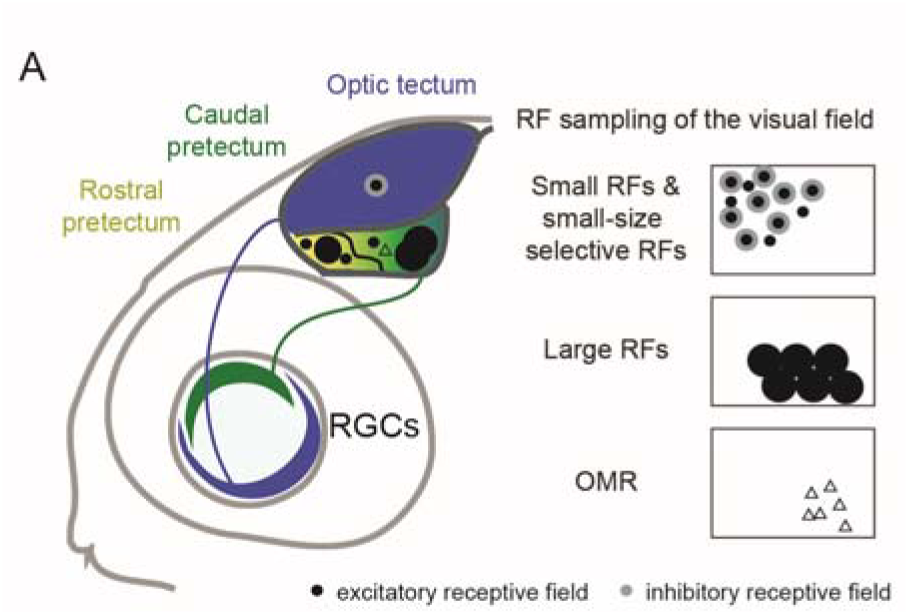
Illustration of the major anatomical locations and visual field preferences of the characterized functional cell types. The tectum mainly contains small-size selective RFs, while a distribution of different RFs is observed in the pretectum, ranging from small-size selective (mainly in the rostral pretectum) to large-size RFs (mainly in the caudal pretectum). Previous findings suggest that tectum and caudal pretectum receive direct retinal inputs (solid lines). To the right, the visual field distributions of RF centres are shown for each functional cell type. An additional visual field map is shown for OMR behaviour, in which the regions driving OMR best are indicated. In the visual field maps, the horizontal axis (left-right) corresponds to the nasal-temporal spatial locations and the vertical axis to the upper and lower visual field locations, respectively.

In summary, we mapped the receptive fields of zebrafish tectal and pretectal neurons and demonstrated that both brain areas fulfil complementary roles for visual motion feature extraction. Receptive fields of tectal neurons are predominantly small, size-selective and have a strong bias in representing the upper-nasal visual field (Figure 5). In contrast, caudal pretectal neurons have predominantly larger receptive fields with receptive field centres preferentially located in the lower view field of the animal, which corresponds to the location of strongest OMR drive in the lower-temporal visual field. Thus, each tectal and pretectal brain region extracts different motion stimulus features and samples distinct visual field regions. We speculate that this anisotropic visual field sampling in tectum and pretectum could represent adaptations of zebrafish to feeding and stabilisation behaviours, resulting in efficient usage of visual brain area volumes for the representation of behaviourally relevant stimulus features. Our study reveals the sensory layout of motion processing and thus constitutes an important advance for deriving a biologically faithful model of visuomotor transformations in zebrafish.

## Author contributions

KW performed the experiments on pretectal and tectal somatic responses. KW, JH and YZ analysed the data. ABA, KW, JH and TRT conceived the experiments and associated analysis protocols. KW, JH, and ABA wrote the manuscript, with input from TRT.

## Acknowledgments

We thank Väinö Haikala and Dierk F. Reiff for help with the visual stimulus arena. The identification of the anatomical position of pretectal dopaminergic neurons was conducted in collaboration with Wolfgang Driever and Christian Altbürger, who provided anatomical z-stacks of a double-transgenic dopaminergic line (unpublished, see (Fernandes et al., 2012; Reinig et al., 2017)). We furthermore thank Thomas Nieß (glassblower shop, University of Tübingen) and Klaus Vollmer (fine mechanics workshop, University Clinic Tübingen) for technical support, Prudenter-Agas (Hamburg, Germany) for generating illustrations, Julianne Skinner for contributing to the analysis of RF centre distribution. We thank Martin Meyer for feedback on a previous version of the manuscript. This work was funded by the Deutsche Forschungsgemeinschaft (DFG) grants EXC307 (CIN – Werner Reichardt Centre for Integrative Neuroscience) and INST 37/967-1 FUGG, and a Human Frontier Science Program (HFSP) Young Investigator Grant RGY0079.

## Quantification and statistical analysis

(In the submitted manuscript, the statistical information is provided in each of the sections above. If our submission is accepted, we will format the Methods section to fully comply with the Star Methods format.)

The analysed number of zebrafish and brains is indicated in the main text and figure legends. Error bars correspond to SEM unless stated otherwise.

## Data and software availability

The scripts for data pre-processing are freely available from our g-node repository (https://web.gin.g-node.org/Arrenberg_Lab/Directional_selectivity_data). All raw and processed data and software used to generate the figures will be made available upon request.

## STAR Methods

### CONTACT FOR REAGENTS AND RESOURCE SHARING

Further information and requests for resources and requests for resources and reagents should be directed to the Lead Contact Aristides Arrenberg.

### EXPERIMENTAL MODEL AND SUBJECT DETAILS

#### Animal care and transgenic lines

All animal procedures conformed to the institutional guidelines of the Universities of Tübingen and Freiburg and the local government (Regierungspräsidium Tübingen and Regierungspräsidium Freiburg, respectively). The transgenic zebrafish line *Tg(HuC:GCaMP5G)a4598Tg* was used in this study. Transgenic lines were kept in either a TL or TLN (nacre) background. Zebrafish larvae were raised in E3 medium until day 5 or 6 post-fertilization (dpf).

### METHOD DETAILS

#### Animal preparation (tectal and pretectal imaging of neuronal somata)

At the day of experiments (5 or 6 dpf), larvae were transferred into a petri dish and embedded in low melting agarose (E3 medium). The agarose surrounding the eyes was not removed as to minimize the range of possible eye movements. 6 animals received an injection of α-bungarotoxin into the caudal vein to paralyse them and prevent eye movements and motion artefacts. 4 animals were recorded without paralysis. In addition, 6 animals were recorded without paralysis in another independent experiment and exclusively used for the analyses presented in the Supplementary Figures S1B-C, S4B and S4F. The animals were then transferred and mounted in agarose on a glass triangle and the fish head protruded the point of the glass triangle, so that the eyes could see through the (agarose and) water clearly. The agarose surrounding the fish head was trimmed on the sides and in front of the animal to reduce the amount of surrounding agarose (Figure S1F). However, the eyes were still covered by agarose to minimize the range of possible eye movements. The glass triangle was held from the back by a 5 mm thin shaft which was fixed to an 8 cm diameter glass bulb (made by a glass blower) filled with E3 medium. The glass bulb resembled a consumer market light bulb (threading of the light bulb/glass bulb shaft at the back of the fish) and a 5 cm diameter hole was cut on the top of the spherical part to allow for approach of the microscope objective onto the fish. From stimulus arena to the fish eye, the light travelled through air, glass (light hit glass roughly orthogonally in the spherical part as to minimize refraction of light rays), water, and finally agarose. The glass bulb was fixed with its shaft (15.5 mm diameter) to the metal holder which allowed for pitch and yaw adjustments (the glass shaft allowed for adjustments in roll). In the first batches of receptive field mapping (n = 6 larvae) experiments, we noticed a problem regarding reflections on the glass bulb on the opposite side of stimulation, resulting in detected neurons on the ipsilateral side of (intended) stimulation in the tectum (which were not plotted in Figure 3). In the second set of recordings (4 larvae), we wrapped a piece of black, half-cylindrical aluminium foil around the objective. The black aluminium foil was then lowered beyond the eye contralateral to the stimulus to prevent this eye from seeing the reflections. In these recordings, only very few ipsilateral tectal neurons were detected (and were not excluded in the anatomical registration figures). Both sets of recordings were included in data analysis, because only few additional neurons were detected due to the reflections in the ipsilateral tectum, suggesting that the vast majority of the detected neurons of the contralateral side were detected due to the stimulus presented to the intended eye.

#### 2-photon microscopy of somatic calcium responses

Calcium imaging was performed with a two-photon microscopy setup based on the MOM microscope (Sutter Instruments, Euler et al. 2009), using a Coherent Vision-S Ti-Sa laser and a 20x/1.0 Zeiss objective to image calcium signals in the transgenic fish line HuC:GCaMP5G (*Tg(elavl3:GCaMP5G)a4598*) (Ahrens et al., 2013b). Calcium time series were recorded at 2 frames per second, with an image size of 512 x 512 pixels and 2 x magnifications, at 920 nm, pre-pulse compensation set to 9756 fs^2^. The midbrain and diencephalon were sampled from +60 µm below the landmark (posterior commissure) to −80 µm above the landmark. Optical slices were taken every 20 µm in the dorso-ventral direction in individual fish and across individual fish, and all dorso-ventral positions were recorded in 10 µm increments relative to the landmark (i.e. no recording at e.g. 5 or 15 µm below the landmark). Since we only recorded every 10 µm in dorso-ventral extent, more than twice as many neurons should have been detectable in the respective brain areas, had we sampled the brain areas at optimal spacing given the neuron soma diameter of ca. 5 µm. Where specified, error bars correspond to measures per completely imaged brain volume. Care was taken to record the same number of slices in each anatomical region. Two animals were used to image one complete brain volume (at 10 µm spacing). Due to the long recording times and positioning instability (likely resulting from the fish drifting within its agarose embedding), we corrected position drifts along the optical axis manually during the recording (mostly less than 4 µm per 30 minutes). Using the 20x Objective and a magnification of 2×, our spatial resolution was 0.43 µm/pixel on the x-axis (medial-lateral) and the y-axis (anterior-posterior).

#### LED arena for visual stimulation (Figures 1-5)

Visual stimulation of zebrafish was conducted with a cylindrical LED arena consisting of 14336 LEDs (Kingbright TA08-81CGKWA): 2 (arena halves) x 8 (rows) x 14 (columns) x 64 (8×8 multiplexed LED matrix) LEDs. The caudal-most column of each arena half was removed without LEDs (i.e. 14, not 15 columns), since the space was needed for the glass bulb stage metal holder. Therefore the caudal-most stimulus patches were slightly cropped (18 ° azimuth instead of 30° azimuth for the last patch). The arena covered −168° to + 168° in azimuth and −40° to 40° in elevation. A few degrees in angle of the dorsal field of view were likely blocked by the objective due to its access angle of 38.39° (<40°), however the eyes were located ∼200 µm below the objective focus which should have resulted in a maximal viewing angle exceeding 38.39° (i.e. 39.2°). The LEDs emitted at 570 nm and an additional high-pass filter foil (LEE no. 779, article 595-1700-7790, castinfo.de, Hagen, Germany) and diffusion filter foil (LEE no. 252, article 595-1780-2520) were placed in front of the arena to optimize GCaMP signal detection and make the stimulus appear more homogeneous. This resulted in a yellow appearance of the stimulus. The LED arena was controlled as described previously (Joesch et al., 2008; Reiser and Dickinson, 2008). LEDs lit during fly-back time of the scanning mirrors.

#### Identification of regions of interest (imaging of neuronal somata)

For analysis of neuronal activity, a custom Matlab script (MOM Load) identified regions based on their correlation to the stimulus and ROIs were manually drawn as described previously (Kubo et al., 2014; Miri et al., 2011). The 3 dimensional mapping of cell location was performed using custom written Matlab scripts (Midbrain_Localizer and Cell_Viewer), which allowed to register the 2 dimensional recordings to a 3 D Z-stack which was acquired after recording sessions (Kubo et al., 2014).

#### 3 D anatomical mapping

To distinguish pretectal from tectal neurons, we proceeded as follows: For each fish, the whole z-stack – which was imaged from the top of tectum to deep ventral pretectum – was resliced to generate a transverse view. On selected, regularly spaced transverse planes (more than ∼50 planes), the ventral border of the tectum was drawn: on each of these transverse plane (512 pixels from left to right, x dimension), a curve was drawn through the area devoid of neuronal somata or fluorescence that was ventrally adjacent to periventricular tectal area with densely packed, fluorescent somata. From each curve, 51 homogeneously distributed points were selected as key points with which a new boundary curve was generated by linear interpolation or three-term Gaussian fitting. Using this method, we obtained a boundary curve with 512 data points corresponding to the pixels in x/y dimensions (left-right and dorsal-ventral) for each transverse plane. In-between the annotated transverse planes, the 2D curves were interpolated to receive a surface that separated the tectum from the pretectum in all three dimensions (Wang et al., 2019). However, the boundaries between the caudal pretectum and adjacent brain areas in the posterior side were not clearly visible in the GCaMP5G fish line. Referring to the AMC structure reported before (Kubo et al., 2014), and to the anatomical annotations of the caudally adjacent tegmentum (Randlett et al., 2015; Ronneberger et al., 2012), the neurons below the tectal-pretectal boundary drawn above, which were located more than 140 um caudally to the posterior commission, were excluded from the pretectum in the current study.

#### Monocular receptive field mapping

In the monocular receptive field mapping experiments, we used horizontally moving gratings, 0.033 cycles/°, moving at 30 °/s, as visual stimuli to induce the neural activities. 8 fish were recorded using naso-temporal motion, and 2 fish were recorded using temporal-nasal motion. The data was pooled because we didn’t observe obvious difference in RF characteristics (RF size, RF centres). The additional recordings for Supplementary Figures S1B, S4B and S4F were performed using both naso-temporal and temporal-nasal motion for 6 fish (3 temporal-nasal and 3 naso-temporal, respectively). Three repetitions of the 57 stimulus phases were shown to the right eye of the fish using one half-cylindrical arena (Figure 1A). All the stimulus patterns were presented in the order depicted in Supplementary Figure S1A. In each trial, every 4.8-s stimulus phase was preceded by a 4 s pause and followed by a 2 s pause with the same stationary visual stimulus pattern. At the beginning and the end of each repetition, we inserted 9 s pauses.

RF maps for individual neurons were calculated by a series of analysis steps. First, we filter the DFF fluorescence traces with a low pass wavelet decomposition [type Daubechies, Matlab: wavedec(DFF,1,’db4’)] and a sliding median filter (the median of three data points). Then deconvolution was performed to the filtered data with the decay time constant (tau) of GCaMP5G, 1.5 s. We calculated the mean of phase-averaged signal (MPAS, averaged over stimulus phase time) from the deconvolved traces. The baseline was defined as the MPAS of all the non-stimulus phases (i.e. without moving stimulus). The standard deviation (STD) of all phase-averaged signals was calculated for the non-motion phases. And the z-score was calculated using the equation [z-score= (MPAS - mean(baseline))/STD(baseline)]. We then calculated the median MPAS z-score (i.e. the median across the three repetitions of a stimulus phase of the average of all data points within one stimulus phase).

To determine the size of receptive fields (RFs) of individual cells, we defined 5 subclasses of responses: small-size receptive fields, medium-size receptive fields, large-size receptive fields, bar-shaped receptive fields and dual receptive fields (containing two discrete excitatory patches in the visual field). Since our stimulation protocol didn’t allow for precise mapping of receptive fields (also Gaussian fits for larger receptive fields were problematic), we turned to a broad classification of RF sizes. To this end we used the smallest stimulation field (30° in azimuth, 13° in elevation) as a calculation unit (i.e. 1 “patch”). After manual inspection of the receptive field locations, we arbitrarily set the thresholds for the 3 size categories as 3 or less active phases (small, SM), 7 or less active phases (medium, ME) and 8 or more active phases (large, L). Bar cells were classified as having active phases only in the full vertical or full horizontal axis, while “double fields” were classified if they had two peaks of activation that were at least 60 ° away from each other. Most double field receptive fields likely result from experimental artefacts, in which stimulus reflections can cause such double field RFs. Double field RFs were excluded from further analysis.

To classify cells according to their respective size criteria, we ran a 2-step process:

***Step I***

1. Calculate mean patch density (MPD)
  a. If the cell responded maximally during the small 6×6 stimulus phases, the MPD corresponded to the average normalized activity of the cell in the phases of the 6×6 stimuli.
  b. If the cell responded maximally for one of the larger stimulus phases, the MPD was calculated as the average normalized activity of the cell in the phases of the 6×6 stimuli that were covered by the field of maximum activation (e.g. the cell in Figure 1D iv had its maximal activity during in the upper horizontal bar stimulus phase, so the mean patch density would correspond to the average activity of the 6 upper patches of the 6×6 stimulation phase).
2. Next, we set the threshold for classifying a part of the visual field as “active” as follows: **IF**
  a. MPD is smaller than 30 %:
    i. MPD * 3
  b. MPD is larger than 30 % and smaller than 40%:
    i. MPD * 2
  c. MPD is larger than 40%:
    i. 90 %
    By relating the activity during the small stimuli to the activity observed during the larger stimuli, these MPD thresholds helped to obtain a more accurate quantification of active patches of cells preferentially active during the small stimulation phases.
3. In *Step I,* cells were classified if following criteria were met:
  a. The maximum activity in the 6×6 stimulation phase exceeded <u>Mean patch density</u> * factor (see step I.2) **or**
  b. Maximum excitation in the 6×6 stimulation phase larger than 90 %
4. To classify the number of active patches, we applied the threshold defined in **point I.2**
  Cells were then either classified as ME or SM based on the number of active patches.

All cells that didn’t meet the criteria from point I.3 were classified according to ***Step II***. First, we calculated a second metric, the mean excitatory density (MED). We calculated the MED by multiplying the calcium response magnitude for each motion phase (excluding the 36 smallest motion phases) with an area factor (full size x 1, half size x 2 …), resulting in a motion phase’s calcium activity weighed by the visual field area in which the stimulus was moving. We call this parameter the “excitatory density” of the stimulated part of the visual field. A biologically plausible underlying cause for differences in excitatory density is the number of DS RGC inputs the cell receives from the portion of the visual field in which the stimulus moves. We then summed the excitatory densities from all larger stimulation fields together (1×1, 2×1, 1×2, 2×2, 1×6, 6×1) taking into account their spatial location in the visual field. This resulted in an excitatory density map of the complete visual field covered by the stimulus arena. In order to report a single number for the RF size, we then defined active phases as those having 75% or more of the normalized summed maximum activation. The difference between this threshold and the 90% threshold for SM cells is derived from comparing manual and automated classification methods.

This method favours smaller receptive fields, because the calcium indicator only shows disproportionally small fluorescence levels for low levels of calcium activity - i.e. it is non-linear - but it enables easy classification of cells size preferences with the given limitations of the calcium indicator (Chen et al., 2013; Pologruto et al., 2004).

The results from our analysis, which is based on thresholding and classification, fits well with both the results obtained from an automated approach using PCA and clustering (Figure S2), and results from manual classification of receptive field sizes.

To determine if cells were small-size selective, we compared the responses to small-size and larger-size stimuli from our stimulus protocol. If (i) a cell was assigned to the small size (SM) or median size (ME) category and was identified during one of the 36 (6×6) small-size stimulation phases (see above), and (ii) the cell showed its maximum activity during the 6×6 stimulation then this cell was classified as being size-selective (‘Inhibition’).

To determine the relative reduction in activity (relative to the response if only the small excitatory receptive field is stimulated) and to visualize the spatial structure of the inhibitory receptive field (“inhibitory density”), we summed the relative reduction of activity (analogous to the above described excitatory density) for every patch belonging to the RF and removed the patches that were part of the RF from the inhibitory field.

Please note that some cells were assigned SM or ME status based on the excitatory density map (and not based on the 36 small-size stimulation phases). We assigned the “no inhibition” status to all of these cells, because the maximal activity was not found in any of the small-size stimulation phases. However, the RFs of these cells could still show some form of inhibition (e.g. during presentation of larger half-field stimulus phases), which was not characterized here. The second cell in Figure 1Dii (an ME cell) is an example for such cells without assigned “inhibition” status.

We estimated the RF centres in XY space as the centre of mass of the normalized activity in the active phases (those with red dots in Figure 1D), i.e. both location and level of activity in active phases determined the position.

The median receptive field size across the visual field was derived by calculating (for every visual field position) the median RF size of receptive fields that covered the respective visual field location to visualize the distribution of RF sizes in the visual space.

LED arena light rays that travelled roughly through the centre of the glass bulb (where the fish head was located as well) hit the glass bulb wall on the opposite side with an angle of incidence of 90°. This resulted in about 4% of reflected light (according to the Fresnel equations) and this light was visible to fish eye that was not intended to be stimulated. In the first 6 animals, we noticed an unexpectedly high number of detected ROIs in the optic tectum ipsilateral to the stimulation. The vast majority of these tectal ipsilateral ROIs had a reversed retino-tectal topography indicating that these ROIs were detected because of the reflected light. We decided to exclude these ROIs in the anatomical reconstruction. For the other 4 animals, we blocked the non-stimulated eye by placing a half-cylindrical piece of black aluminium foil around the objective and lowering it below the level of the eye. In these animals, a much smaller number of neurons were detected in the ipsilateral tectum, and their anatomical location corresponded to the expected retino-tectal topography. In the data analysis, the small-size RF neurons in the ipsilateral hemisphere recorded from the first 6 animals were excluded. The laterality index for tectal neurons was −0.88 for the 4 animals with one blocked eye, and −0.46 for the 6 animals in which the eyes were not blocked.

In Figure 3A (locations of RF centres in the visual field for tectal small-size RF cells), only very few cells (55 out of 1074), which have RFs centred in the lower temporal of the visual field, were present. While we are convinced that this finding represents an actual under-representation of such cells in the optic tectum, we would like to discuss two experimental caveats, which can explain the effect partially (but not fully). First, the fish were mounted on a glass triangle and care was taken to allow free view of the stimuli from the position of the eye lenses by pushing the larva towards the tip of the glass triangle (thereby reducing the positional stability of the recording).

However, for some animals, a small portion of the lower temporal visual field might have been blocked by the sides of the glass triangle. However, this caveat should only have affected extreme lower-temporal receptive field positions (e.g. >120° in azimuth and <-30° in elevation). Second, the anatomical positions of those tectal neurons having lower temporal receptive field centres lie close to the border to the pretectum. The pretectum-classified small-size RF cells in proximity of the tectum (shade red dots in Figure 2D) can fill the gap in the ventral-caudal visual field in Figure 3A only partially (just 12 additional cells for the region > 90° azimuth and < 0° elevation) when such pretectal neurons are plotted together with the tectal neurons.

To characterize the distributions of RF centres across visual space, the density of the RF centres in the visual space was calculated with ‘ksdensity’ function in Matlab. Contour lines were plotted based on the density.

One gap between the rostral and caudal pretectal neurons is quite obvious (Figure 2D). Moreover, in the rostral-ventral pretectum, many neurons are small-size selective (RFs with signs of inhibition). Therefore, a boundary was manually defined between the rostral and caudal pretectum (Figure 2D). In the data analysis of the rostral pretectal neurons, the rostral pretectum was defined as the region (D-V axis < 10 µm and A-P axis < 60 µm, or D-V axis < −30 µm and A-P axis < 110 µm). 140 um caudal to the posterior commissure along the anterior-posterior axis was conceded as the caudal boundary of the pretectum with other brain areas. The bilaterally symmetric anatomical distribution of the neurons relative to the midline was measured using a laterality index, which was calculated as, (neurons_right_ - neurons_left_)/(neurons_left_ + neurons_right_).

#### Inclusion criteria for somatic calcium responses

We calculated the Pearson’s linear correlation coefficients between the stimulus phase z-scores (see above) of the three stimulus protocol repetitions (for all 3 pairwise combinations) to characterize the reproducibility of stimulus-evoked calcium responses. In our further data analysis, we only kept the neurons for which all three correlation coefficients were higher than a certain threshold. The threshold was set between 0.65 and 0.75 to exclude around 30 % neurons with low reproducibility of stimulus-evoked activity in the monocular receptive field mapping experiment (2004 out of 2995, 67% neurons were kept). We then performed signal-to-noise ratio (SNR) analysis to exclude neurons with unstable baseline. In the SNR analysis, a threshold of four was used on the z-score to detect positive neural responses. SNR was defined as the ratio of the average response of all the responsive phases to the standard deviation of the baseline. All neurons with SNR lower than a certain threshold were excluded. The threshold was set between 8 and 10 to exclude about 5% remaining neurons with low SNR. We kept about 96% (1926 out of 2004) in the monocular receptive field mapping experiment for further analysis.

#### Setup for measuring the receptive field of OMR behaviour

The visual stimuli were presented binocularly with a 336° (from −168° to 168° in azimuth) surround LED arena (two half-cylindrical arenas, Figure 4A) from both sides of the fish, i.e. all stimulus phases (except the rotational and looming control stimuli) were mirror-symmetric across the midline. The image of the fish was reflected to a lens by a mirror positioned around 5 cm above the fish (1 cm above the glass bulb). An infrared-sensitive high speed camera (Modell IDT iNdustrial Speed I, Integrated Design Tools Inc.) with an IR bandpass filter (ET 850/40, CHROMA) recorded (250 Hz) the behaviours of the fish through the lens (diameter, 25.4 mm; focal length, 100mm; THORLABS, LB1676) mentioned above. Since the mirror above the fish is tilted 45 degrees vertically, the light path from the mirror to the camera was horizontal. The fish was illuminated from below with a high power infrared LED light (850 nm, Conrad, Item No. 491248-62) positioned 1 cm below the glass bulb. The infrared light was mounted below a piece of milk glass and provided homogenous background illumination around the fish (Figure S5A). The infrared LED light and the camera were triggered by the motion signal of the visual stimulus recorded by a LabVIEW DAQ box. In our experiment, the camera started recording about 300 milliseconds after the motion phase onset of the visual stimuli. The infrared LED light was only on during the motion visual stimulus phases to reduce potential harm to the fish.

#### Receptive field mapping of OMR behaviour

For each recording, a larval zebrafish (6 dpf) was transferred into a petri dish and embedded in low melting agarose (E3 medium) on a glass triangle such that the fish body completely protruded the tip of the glass triangle. The agarose surrounding the larval tail was removed to free the tail. To reduce the amount of agarose surrounding the animal, agarose surrounding the fish was trimmed on the sides and in front. Only the agarose caudal to the animal attached to the triangular stage. Therefore, a large view field was accessible for the fish, and - more importantly - the contours of the beating tail could be imaged without optical obstruction from the glass triangle. The fish and the glass triangle were fixed to an 8 cm diameter glass bulb filled with E3 medium in the same way as described above (see Animal preparation in the Method section). The animal was illuminated from below with a high power infrared LED light, which was triggered on 150 milliseconds after the start of the grating motion and lasted for 10 s. During the first 6 s of each grating motion phase (300 ms delay), the animal was imaged with a high speed infrared camera at 250 frames per second. The video data were saved during the pauses and analysed with custom written Matlab code offline.

In the OMR behavioural test, forward moving gratings, 0.033 cycles/°, moving at 30°/s, were used as visual stimuli to induce the optomotor response. Three repetitions of the 55 stimulus phases were shown to both eyes of the fish using two half-cylindrical arenas. All the stimulus patterns were presented in each of the three randomized orders depicted in Supplementary Figure S6 for the repetitions. The animals were adapted to the stationary gratings for about half an hour before motion stimulation started. In each trial, every 9.75-s stimulus phase was preceded by an 18 s pause and followed by a 51 s pause with the same stationary visual stimulus pattern. Before presenting the 2^nd^ and 3^rd^ repetition of the 55 stimulus phases, the animals rested for about 1 hour.

The motion of the larval tail was traced using a custom Matlab algorithm for image processing and the ‘alpha angle’ (the angle between the fish anterior-posterior body axis and the tail tip, see Figure 4A) was calculated. The single tail beats were detected by labelling the peaks and troughs of the alpha angle traces. In our analysis, only swim beats meeting the following two criteria were considered as forward OMR beats: (1) the tail-beat frequency (during individual swimming bouts) was higher than 25 Hz; (2) for each beat, the difference of the amplitudes of adjacent peaks and troughs divided by the sum of them was smaller than 0.3 and larger than −0.3 (symmetrical tail beats/forward swimming).

Two types of OMR tail beats were distinguished: symmetrical and unsymmetrical tail beats (Figures 4E and S5E). Unsymmetrical tail beats oftentimes occurred at the beginning and the end of OMR tail beat bouts (Figures 4E and S5E), similar to the case of freely swimming larval zebrafish (Marques et al., 2018). As expected, the larval zebrafish tried to turn in response to the rotating visual stimulus used as control in our protocol, with tails beating mainly unilaterally to the reverse direction of the rotation (Ahrens et al., 2013a). However, symmetrical tail beats were observed in the turning swimming bouts as well (Figure S5Ei).

In Figure 4 (and Supplementary Figure S5), the shown tail-beat frequency does not correspond to the frequency reached during individual swim bouts, but instead corresponds to the average frequency during the stimulus phase, such that time periods without swim bouts lead to a reduction of tail-beat frequency.

## Supplementary Figures

**Supplementary Figure S1 (related to Figures 1 and 2).**
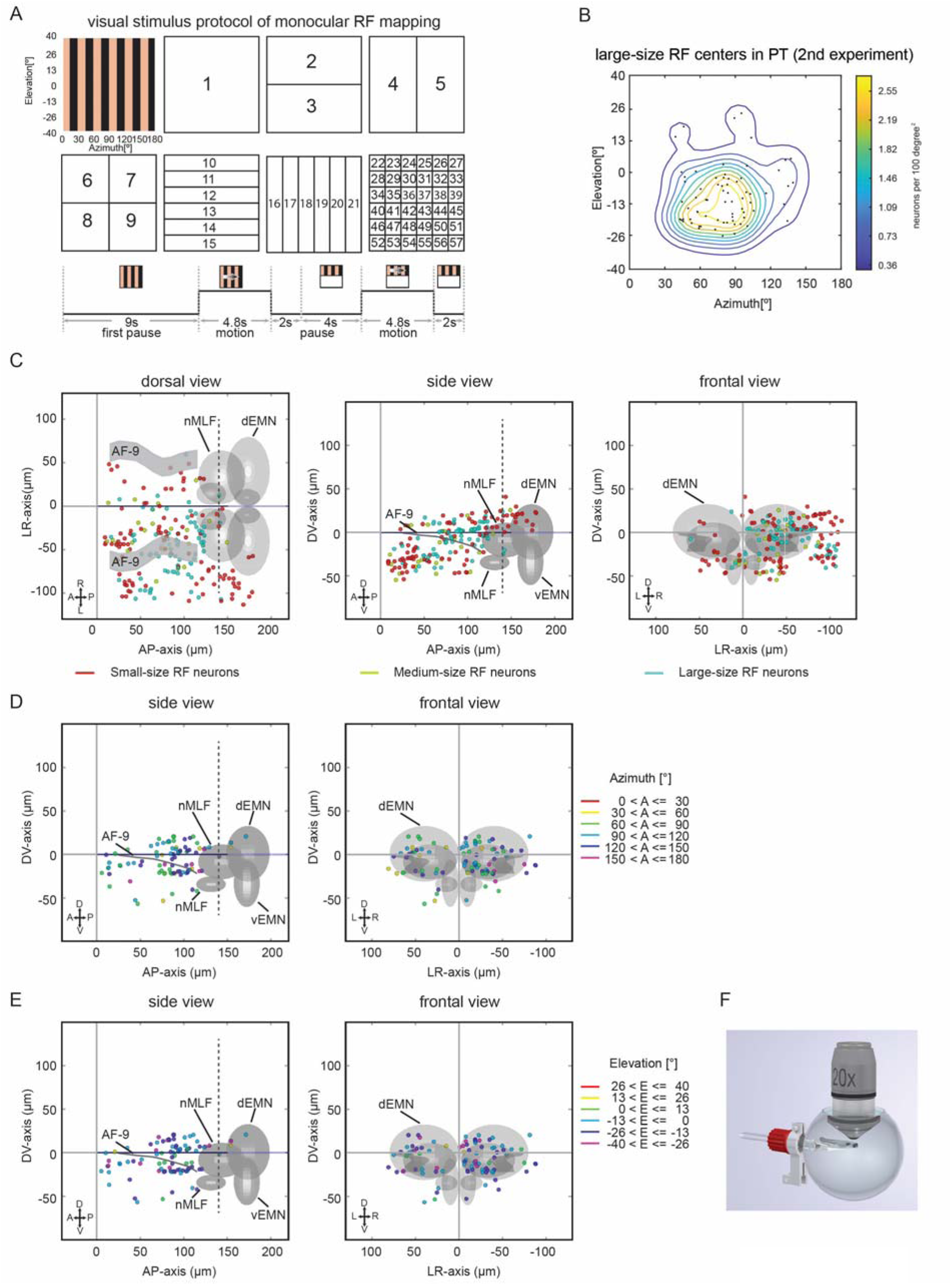
Stimulus protocols for calcium imaging of neuronal somata and distribution of large-size RF pretectal neurons. **(A)** Stimulus protocol for the monocular receptive field mapping experiment. Upper: Each rectangle corresponds to the stimulus phases shown in Figure 1A. The coordinate system of the visual field is indicated in the first plot with gratings. The numbers in the other plots indicate the temporal order of motion stimuli within the stimulus protocol. The 57 motion patterns were presented in three repetitions in each recording. Bottom: Each motion phase, lasting 4.8 s, was preceded by a pause phase of 6 s, in which stationary gratings were shown. **(B)** Locations and density contour plot of receptive field centres of pretectal large-size RF neurons (n = 6 pretecta; 2^nd^ experiment/new dataset). **(C)** Anatomical maps of small-size (red), medium-size (yellow) and large-size RF (cyan) neurons in the pretectal region of the second newly recorded dataset (n = 6 pretecta). **(D, E)** Topographic maps of large-size RF neurons in the pretectal region. Each coloured dot represents a single neuron with its receptive field centre in the corresponding azimuth **(D)** and elevation **(E)** range. For example, all receptive field centres of the neurons in red are located in-between 0° azimuth (in front of the fish) and 30° azimuth (to the right side of the fish) in panel **(D)**. All receptive field centres of the neurons in green are located in-between 0° in elevation (equator of the view field) and 13° elevation (upper view field) in panel (E). **(F)** The larval zebrafish were embedded in low melting agarose on a small triangle stage located in a glass bulb (the schematic is not drawn to scale; the diameter of the glass bulb was approximately 8 cm).

**Supplementary Figure S2.**
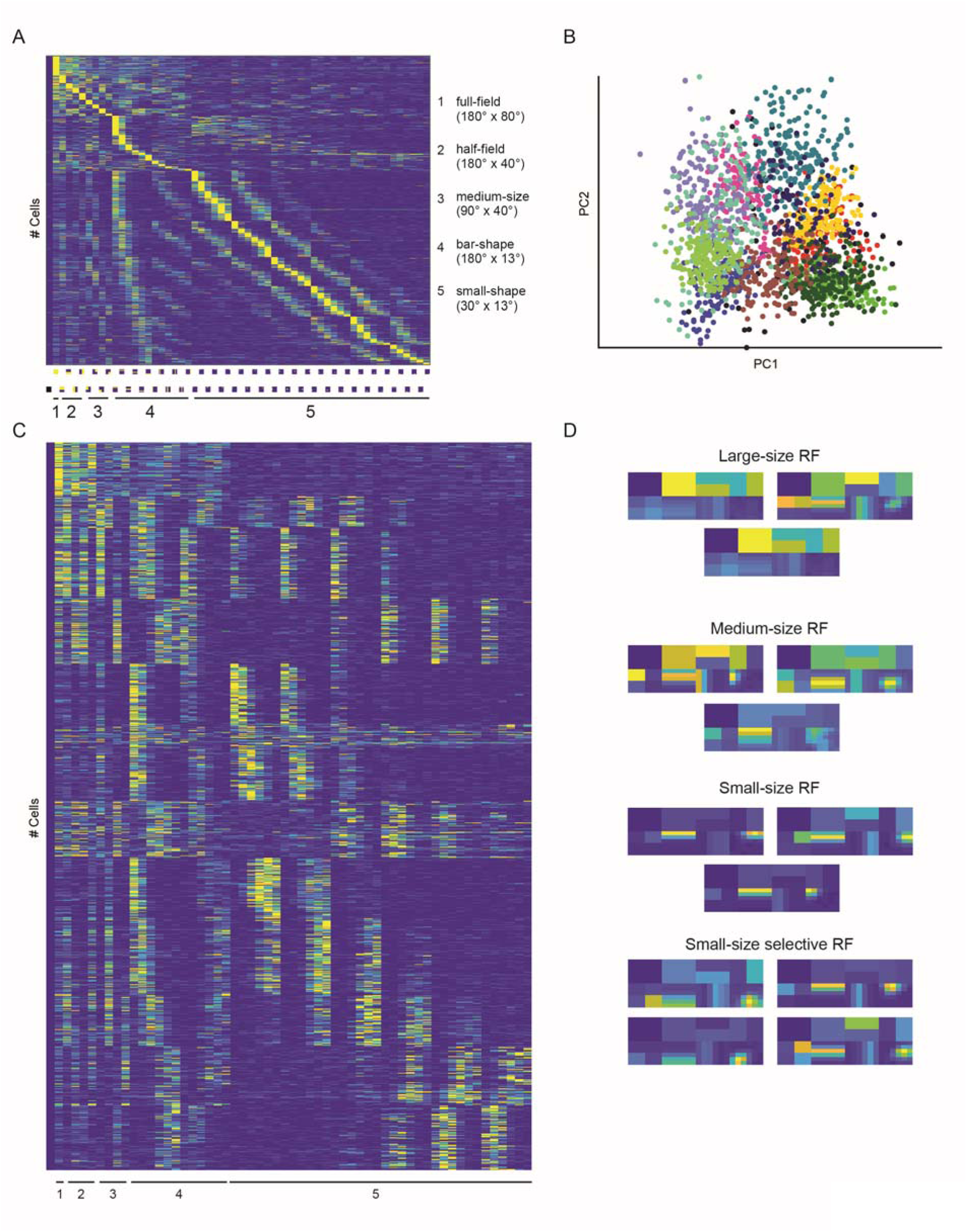
Unbiased classification of receptive field sizes confirms the presented results, which are based on thresholding. **(A)** Cellular responses towards all 57 stimulus phases for all recorded cells, sorted according to the stimulation phase with the maximal response. **(B)** Result of principal component (PC) analysis and expectation maximization clustering, splitting the cell responses in 13 distinct response groups. **(C)** The cellular responses sorted according to their respective cluster and the stimulus phase with its maximal response (from large to small). **(D)** Average responses of all cells belonging to a particular response group during all 57 stimulus phases. We split them according to their receptive field size and into the main categories shown in the main figure.

**Supplementary Figure S3 (related to Figure 3).**
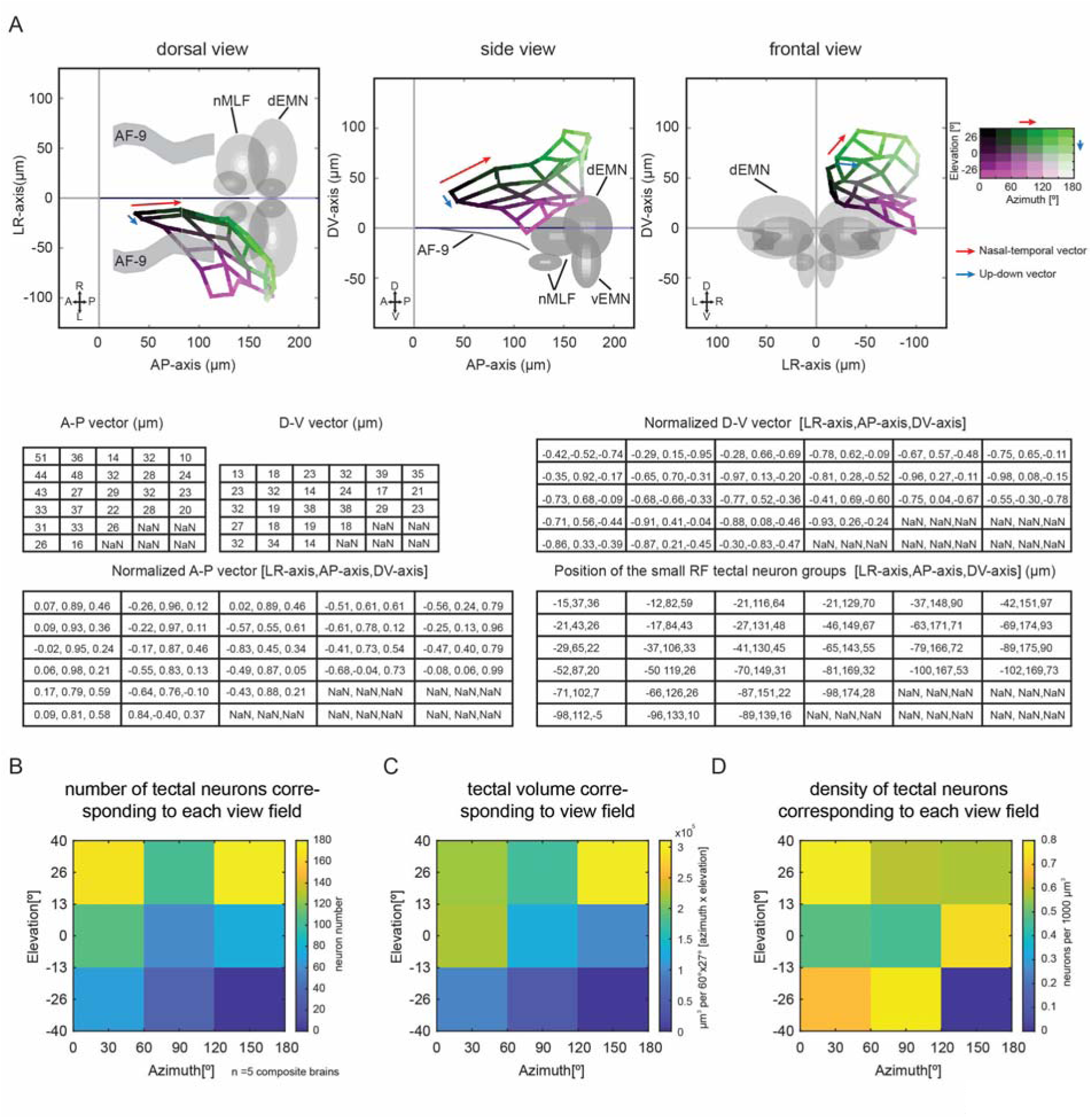
**(A)** Upper: anatomical map of the tectal receptive field centre topography (Figure 3B). The average anatomical position of neurons representing different visual field bins was calculated based on the RF centres in each of the 6 x 6 visual field bins. All 36 locations were connected by a grid to illustrate the mapping of visual space (legend on the right) to anatomical space in the tectum. Examples of nasal-temporal (N-T) and up-down (U-D) vectors are illustrated with red and blue arrows. Below: tables of the lengths and directions of the N-T and U-D vectors in 3D and median positions of all the small-size RF tectal neuronal bins with RF centres in the corresponding 6 x 6 view fields. **(B)** The number of small-size RF tectal neurons with their receptive filed centres in the corresponding 3 x 3 areas of the whole view field. **(C)** The brain volumes enclosing the small-size RF tectal neurons with their receptive filed centres in the corresponding 3 x 3 areas of the whole view field. **(D)** The density of small-size RF tectal neurons with their receptive filed centres in the corresponding 3 x 3 areas of the whole view field.

**Supplementary Figure S4 (related to Figure 2).**
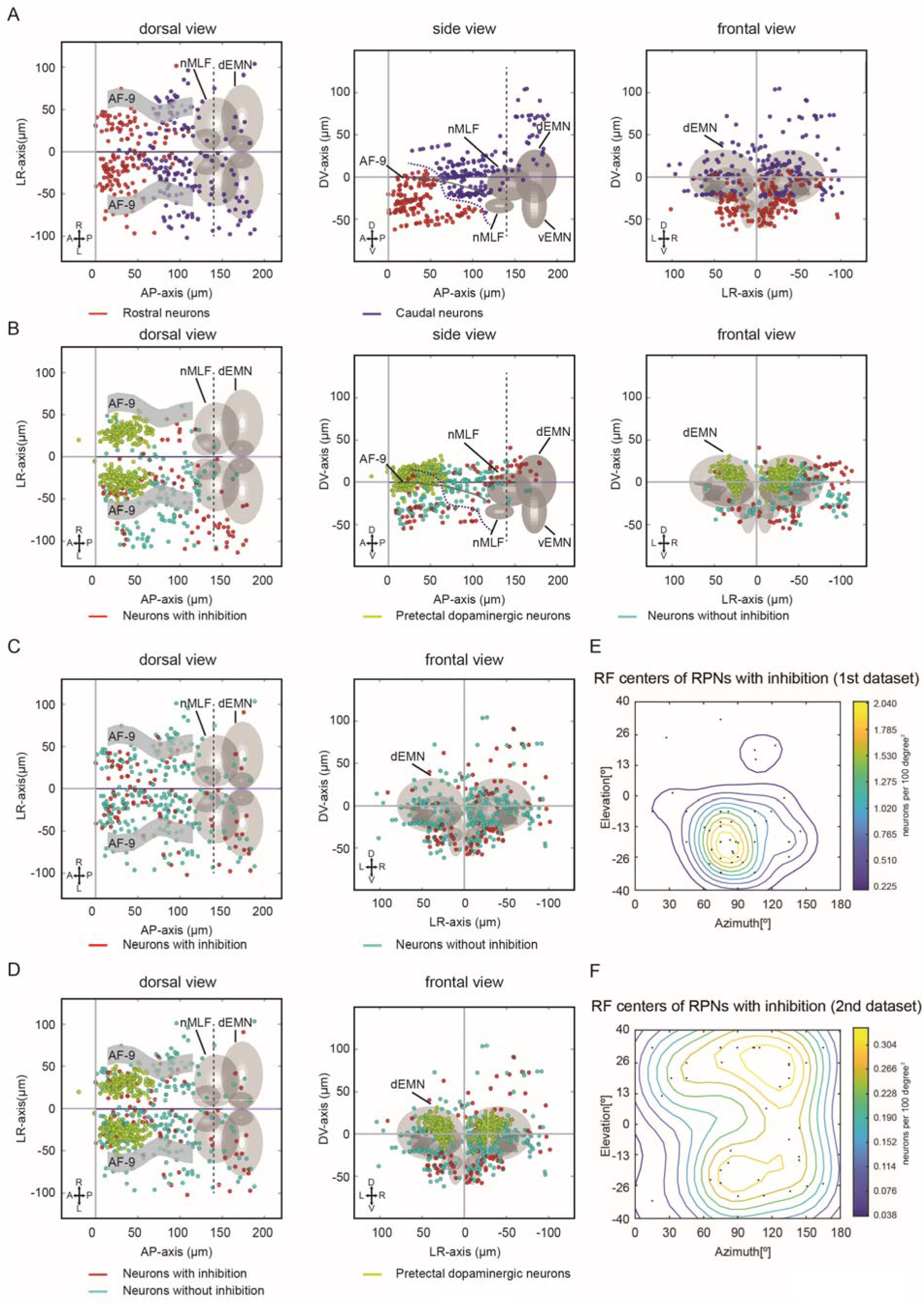
**(A)** Anatomical map of neurons in the rostral (red) and caudal (blue) pretectal regions. Each coloured dot represents a single neuron. The dashed blue curve in the centre panel shows the boundary between the rostral and caudal pretectum (n = 10 fish, 5 composite pretectal regions). **(B)** Anatomical map of three types of neurons in the pretectal region: neurons with (red)/without (cyan) inhibition and pretectal dopaminergic neurons (green) from the 2^nd^ dataset (n = 6 fish, 6 pretecta). Each coloured dot represents a single neuron. **(C)** Anatomical map of two types of neurons in the pretectal region (neurons with (red)/without (cyan) inhibition) from the 1^st^ dataset. Each coloured dot represents a single neuron (n = 10 fish, 5 composite pretectal regions). **(D)** Anatomical map of three types of neurons in the pretectal region from the 1^st^ dataset: neurons with (red)/without (cyan) inhibition and pretectal dopaminergic neurons (green). Each coloured dot represents a single neuron (n = 10 fish, 5 composite pretectal regions). Coordinates are defined as distances relative to the posterior commissure in the diencephalon (anterior-posterior axis and dorso-ventral axis) and midline (left-right axis). **(E)** Locations and density contour plot of receptive field centres of rostral pretectal neurons (RPNs) with inhibition in the 1^st^ dataset (n = 10 fish, 5 composite pretectal regions). **(F)** Locations and density contour plot of receptive field centres of rostral pretectal neurons with inhibition in the 2^nd^ dataset (n = 6 fish, 6 pretecta).

**Supplementary Figure S5 (related to Figure 4).**
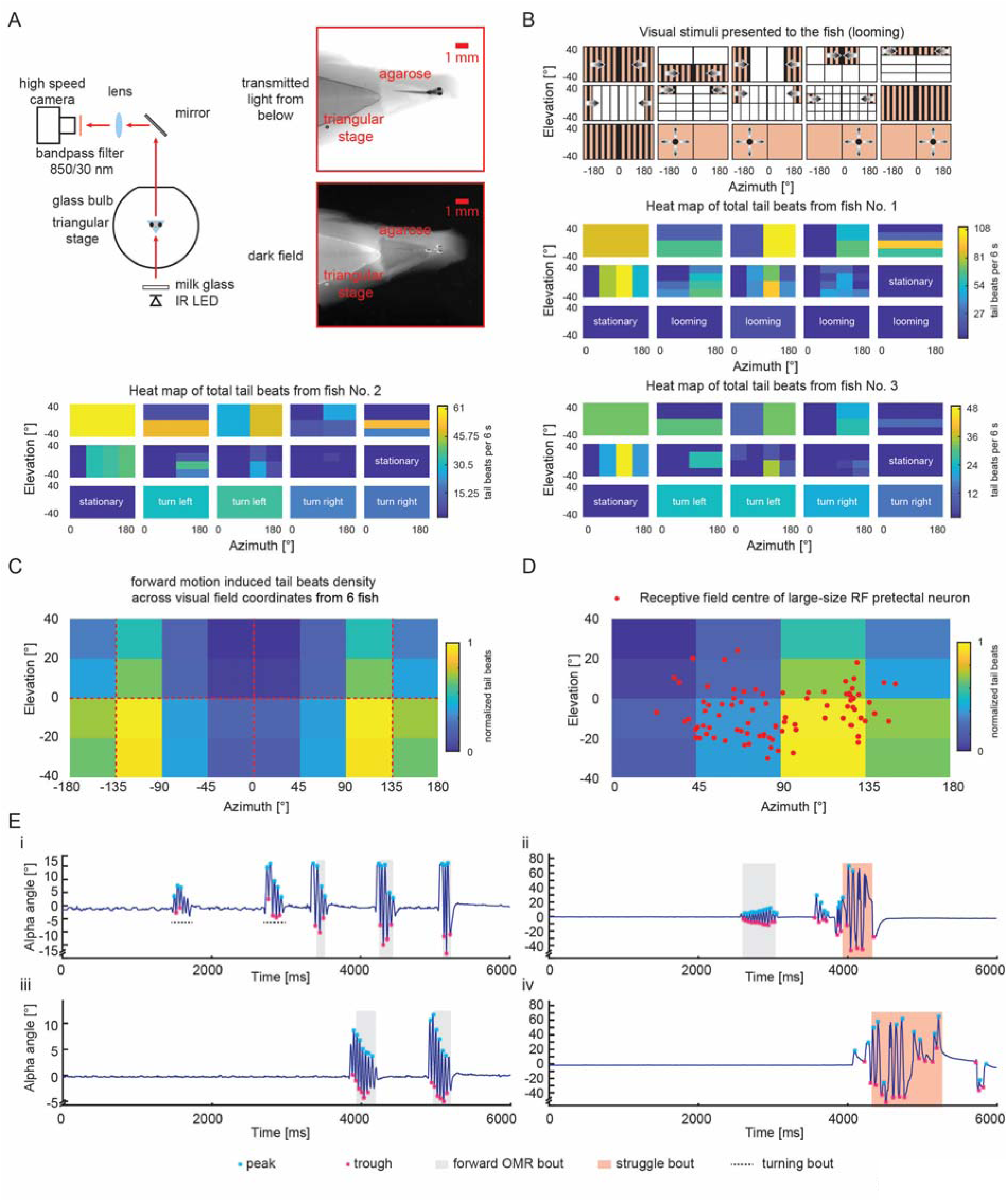
**(A)** Frontal view of the schematic of the setup for the OMR behavioural test (Cylindrical arena for presenting visual stimulus is not shown). A photo of an embedded larval zebrafish with the tail free for behaviour test (right). **(B)** Illustration of the visual stimulus patterns (the protocol with looming, only for fish no. 1) and heat maps of the total tail beat count (including unsymmetrical tail beats) medians induced by visual stimulus for three larval zebrafish. In the last 4 phases, looming visual stimuli were shown on either side to fish no. 1 instead of rotational binocular whole field motion (Supplementary Figure S6). The non-stimulated regions are shown as white areas for illustration purposes but contained a stationary grating. **(C)** Heat maps of the tail movement density across visual field coordinates (n = 6; see Method). Here all tail beats are included, whereas in Figure 4, forward-OMR-like tail beats (symmetrical tail beats) were selectively included. **(D)** RF centres of the large-size RF pretectal neurons (red dots; also see Figure 2C) plotted on the heat maps of the OMR density across visual field coordinates from six fish. **(E)** Tail movements induced by rotational (i) and forward (ii-iv) moving gratings of different sizes and locations. The peaks and troughs of each swim bout within the 6-s recordings are labelled with cyan and magenta asterisks. OMR forward swimming beats and struggling tail beats are indicated with grey and pink background shades, respectively. Identified asymmetric (turning) tail beats are labelled by a dashed line.

**Supplementary Figure S6 (related to Figure 4).**
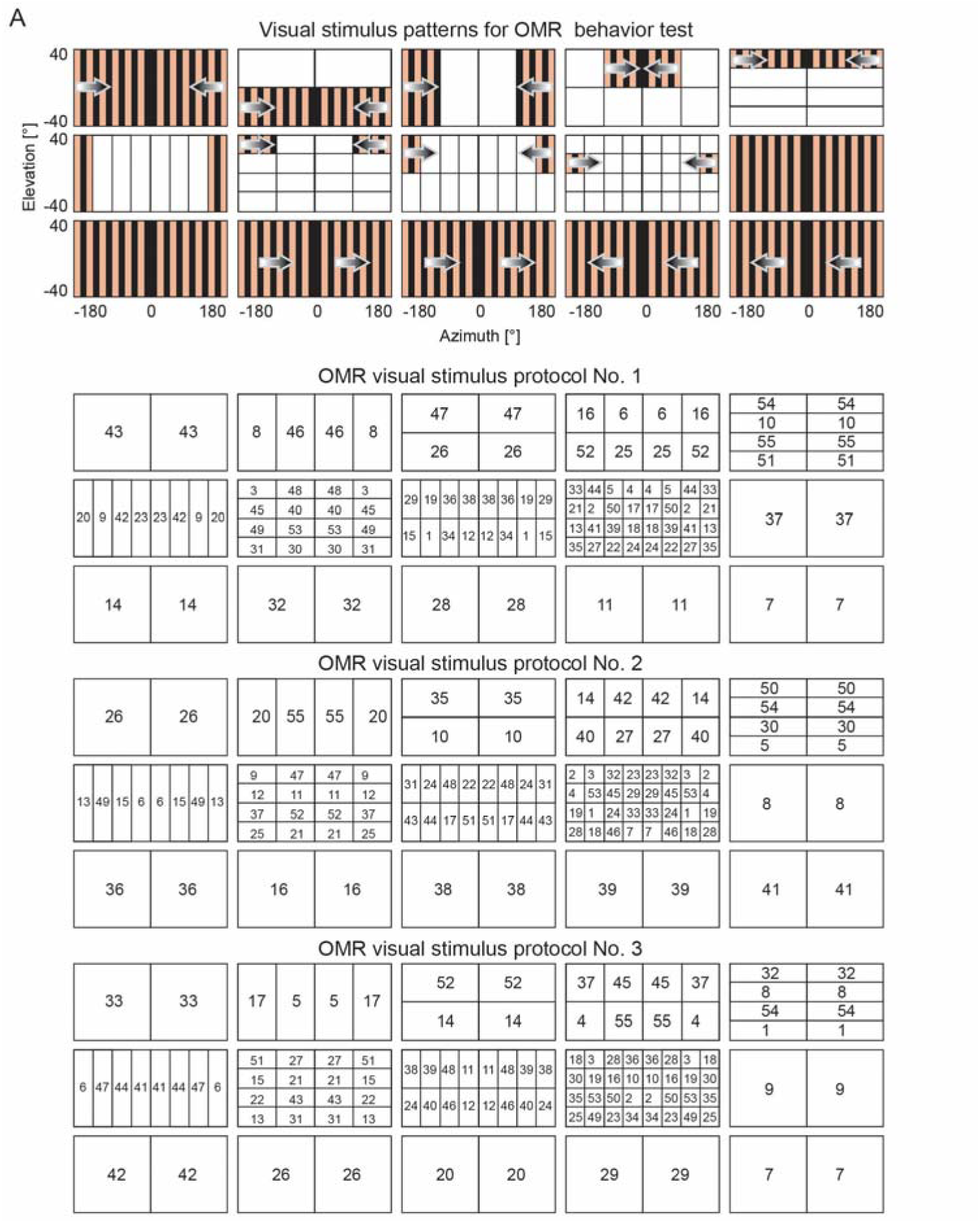
Three visual stimulus protocols for the OMR behaviour test. Upper panel: motion stimuli of the OMR experiment (the whole protocol shown in truncated form in Figure 4B) as a reference for the protocols shown below. Lower panels: the numbers indicate the order in which the visual stimuli were presented to the animals in each protocol. All three protocols were presented to each animal in sequence, resulting in three repetitions of each stimulus phase.

**Supplementary Movie S1. OMR behaviour induced by whole field forward moving gratings from both sides.** OMR swimming bouts were recorded at 250 frames per second (fps) for 6 seconds. The broadcasting frame rate was reduced to 60 fps (0.24× speed).

## References

Ahrens, M.B., Huang, K.H., Narayan, S., Mensh, B.D., and Engert, F. (2013a). Two-photon calcium imaging during fictive navigation in virtual environments. Front Neural Circuits 7, 104.

Ahrens, M.B., Orger, M.B., Robson, D.N., Li, J.M., and Keller, P.J. (2013b). Whole-brain functional imaging at cellular resolution using light-sheet microscopy. Nat Methods 10, 413–420.

Arrenberg, A.B., and Driever, W. (2013). Integrating anatomy and function for zebrafish circuit analysis. Front Neural Circuits 7, 74.

Arunachalam, M., Raja, M., Vijayakumar, C., Malaiammal, P., and Mayden, R.L. (2013). Natural history of zebrafish (Danio rerio) in India. Zebrafish 10, 1–14.

Attardi, D.G., and Sperry, R.W. (1963). Preferential selection of central pathways by regenerating optic fibers. Exp Neurol 7, 46–64.

Baden, T., Berens, P., Franke, K., Roman Roson, M., Bethge, M., and Euler, T. (2016). The functional diversity of retinal ganglion cells in the mouse. Nature 529, 345–350.

Baier, H., Klostermann, S., Trowe, T., Karlstrom, R.O., Nusslein-Volhard, C., and Bonhoeffer, F. (1996). Genetic dissection of the retinotectal projection. Development 123, 415–425.

Beck, J.C., Gilland, E., Tank, D.W., and Baker, R. (2004). Quantifying the ontogeny of optokinetic and vestibuloocular behaviors in zebrafish, medaka, and goldfish. J Neurophysiol 92, 3546–3561.

Bergmann, K., Meza Santoscoy, P., Lygdas, K., Nikolaeva, Y., MacDonald, R.B., Cunliffe, V.T., and Nikolaev, A. (2018). Imaging Neuronal Activity in the Optic Tectum of Late Stage Larval Zebrafish. J Dev Biol 6.

Bianco, I.H., Kampff, A.R., and Engert, F. (2011). Prey capture behavior evoked by simple visual stimuli in larval zebrafish. Front Syst Neurosci 5, 101.

Boulanger-Weill, J., Candat, V., Jouary, A., Romano, S.A., Perez-Schuster, V., and Sumbre, G. (2017). Functional Interactions between Newborn and Mature Neurons Leading to Integration into Established Neuronal Circuits. Curr Biol 27, 1707–1720 e1705.

Britto, L.R., Natal, C.L., and Marcondes, A.M. (1981). The accessory optic system in pigeons: receptive field properties of identified neurons. Brain Res 206, 149–154.

Busch, C., Borst, A., and Mauss, A.S. (2018). Bi-directional Control of Walking Behavior by Horizontal Optic Flow Sensors. Curr Biol 28, 4037–4045 e4035.

Chen, T.W., Wardill, T.J., Sun, Y., Pulver, S.R., Renninger, S.L., Baohan, A., Schreiter, E.R., Kerr, R.A., Orger, M.B., Jayaraman, V., et al. (2013). Ultrasensitive fluorescent proteins for imaging neuronal activity. Nature 499, 295–300.

Cowey, A., and Rolls, E.T. (1974). Human cortical magnification factor and its relation to visual acuity. Exp Brain Res 21, 447–454.

Fernald, R.D., and Shelton, L.C. (1985). The organization of the diencephalon and the pretectum in the cichlid fish, Haplochromis burtoni. J Comp Neurol 238, 202–217.

Fernandes, A.M., Fero, K., Arrenberg, A.B., Bergeron, S.A., Driever, W., and Burgess, H.A. (2012). Deep brain photoreceptors control light-seeking behavior in zebrafish larvae. Curr Biol 22, 2042–2047.

Filippi, A., Mueller, T., and Driever, W. (2014). vglut2 and gad expression reveal distinct patterns of dual GABAergic versus glutamatergic cotransmitter phenotypes of dopaminergic and noradrenergic neurons in the zebrafish brain. J Comp Neurol 522, 2019–2037.

Gahtan, E., Tanger, P., and Baier, H. (2005). Visual prey capture in larval zebrafish is controlled by identified reticulospinal neurons downstream of the tectum. J Neurosci 25, 9294–9303.

Giolli, R.A., Blanks, R.H., and Lui, F. (2006). The accessory optic system: basic organization with an update on connectivity, neurochemistry, and function. Prog Brain Res 151, 407–440.

Grama, A., and Engert, F. (2012). Direction selectivity in the larval zebrafish tectum is mediated by asymmetric inhibition. Front Neural Circuits 6, 59.

Grasse, K.L., and Cynader, M.S. (1984). Electrophysiology of lateral and dorsal terminal nuclei of the cat accessory optic system. J Neurophysiol 51, 276–293.

Grujic, N., Brehm, N., Gloge, C., Zhuo, W., and Hafed, Z.M. (2018). Perisaccadic perceptual mislocalization is different for upward saccades. J Neurophysiol 120, 3198–3216.

Hafed, Z.M., and Chen, C.Y. (2016). Sharper, Stronger, Faster Upper Visual Field Representation in Primate Superior Colliculus. Curr Biol 26, 1647–1658.

Haikala, V., Joesch, M., Borst, A., and Mauss, A.S. (2013). Optogenetic control of fly optomotor responses. J Neurosci 33, 13927–13934.

Hunter, P.R., Lowe, A.S., Thompson, I.D., and Meyer, M.P. (2013). Emergent properties of the optic tectum revealed by population analysis of direction and orientation selectivity. J Neurosci 33, 13940–13945.

Joesch, M., Plett, J., Borst, A., and Reiff, D.F. (2008). Response properties of motion-sensitive visual interneurons in the lobula plate of Drosophila melanogaster. Curr Biol 18, 368–374.

Karmeier, K., Krapp, H.G., and Egelhaaf, M. (2003). Robustness of the tuning of fly visual interneurons to rotatory optic flow. J Neurophysiol 90, 1626–1634.

Kastenhuber, E., Kratochwil, C.F., Ryu, S., Schweitzer, J., and Driever, W. (2010). Genetic dissection of dopaminergic and noradrenergic contributions to catecholaminergic tracts in early larval zebrafish. J Comp Neurol 518, 439–458.

Krapp, H.G., Hengstenberg, R., and Egelhaaf, M. (2001). Binocular contributions to optic flow processing in the fly visual system. J Neurophysiol 85, 724–734.

Kubo, F., Hablitzel, B., Dal Maschio, M., Driever, W., Baier, H., and Arrenberg, A.B. (2014). Functional architecture of an optic flow-responsive area that drives horizontal eye movements in zebrafish. Neuron 81, 1344–1359.

Marques, J.C., Lackner, S., Felix, R., and Orger, M.B. (2018). Structure of the Zebrafish Locomotor Repertoire Revealed with Unsupervised Behavioral Clustering. Curr Biol 28, 181–195 e185.

Masseck, O.A., and Hoffmann, K.P. (2008). Responses to moving visual stimuli in pretectal neurons of the small-spotted dogfish (Scyliorhinus canicula). J Neurophysiol 99, 200–207.

Miri, A., Daie, K., Burdine, R.D., Aksay, E., and Tank, D.W. (2011). Regression-based identification of behavior-encoding neurons during large-scale optical imaging of neural activity at cellular resolution. J Neurophysiol 105, 964–980.

Muto, A., Lal, P., Ailani, D., Abe, G., Itoh, M., and Kawakami, K. (2017). Activation of the hypothalamic feeding centre upon visual prey detection. Nat Commun 8, 15029.

Nassi, J.J., and Callaway, E.M. (2009). Parallel processing strategies of the primate visual system. Nat Rev Neurosci 10, 360–372.

Naumann, E.A., Fitzgerald, J.E., Dunn, T.W., Rihel, J., Sompolinsky, H., and Engert, F. (2016). From Whole-Brain Data to Functional Circuit Models: The Zebrafish Optomotor Response. Cell 167, 947–960 e920.

Niell, C.M., and Smith, S.J. (2005). Functional imaging reveals rapid development of visual response properties in the zebrafish tectum. Neuron 45, 941–951.

Orger, M.B., Kampff, A.R., Severi, K.E., Bollmann, J.H., and Engert, F. (2008). Control of visually guided behavior by distinct populations of spinal projection neurons. Nat Neurosci 11, 327–333.

Pologruto, T.A., Yasuda, R., and Svoboda, K. (2004). Monitoring neural activity and [Ca2+] with genetically encoded Ca2+ indicators. J Neurosci 24, 9572–9579.

Portugues, R., and Engert, F. (2009). The neural basis of visual behaviors in the larval zebrafish. Curr Opin Neurobiol 19, 644–647.

Presson, J., Fernald, R.D., and Max, M. (1985). The organization of retinal projections to the diencephalon and pretectum in the cichlid fish, Haplochromis burtoni. J Comp Neurol 235, 360–374.

Preuss, S.J., Trivedi, C.A., vom Berg-Maurer, C.M., Ryu, S., and Bollmann, J.H. (2014). Classification of object size in retinotectal microcircuits. Curr Biol 24, 2376–2385.

Ramdya, P., and Engert, F. (2008). Emergence of binocular functional properties in a monocular neural circuit. Nat Neurosci 11, 1083–1090.

Randlett, O., Wee, C.L., Naumann, E.A., Nnaemeka, O., Schoppik, D., Fitzgerald, J.E., Portugues, R., Lacoste, A.M., Riegler, C., Engert, F., et al. (2015). Whole-brain activity mapping onto a zebrafish brain atlas. Nat Methods 12, 1039–1046.

Recher, G., Jouralet, J., Brombin, A., Heuze, A., Mugniery, E., Hermel, J.M., Desnoulez, S., Savy, T., Herbomel, P., Bourrat, F., et al. (2013). Zebrafish midbrain slow-amplifying progenitors exhibit high levels of transcripts for nucleotide and ribosome biogenesis. Development 140, 4860–4869.

Reinig, S., Driever, W., and Arrenberg, A.B. (2017). The Descending Diencephalic Dopamine System Is Tuned to Sensory Stimuli. Curr Biol 27, 318–333.

Reiser, M.B., and Dickinson, M.H. (2008). A modular display system for insect behavioral neuroscience. J Neurosci Methods 167, 127–139.

Rinner, O., Rick, J.M., and Neuhauss, S.C. (2005). Contrast sensitivity, spatial and temporal tuning of the larval zebrafish optokinetic response. Invest Ophthalmol Vis Sci 46, 137–142.

Robles, E., Laurell, E., and Baier, H. (2014). The retinal projectome reveals brain-area-specific visual representations generated by ganglion cell diversity. Curr Biol 24, 2085–2096.

Romano, S.A., Pietri, T., Perez-Schuster, V., Jouary, A., Haudrechy, M., and Sumbre, G. (2015). Spontaneous neuronal network dynamics reveal circuit’s functional adaptations for behavior. Neuron 85, 1070–1085.

Ronneberger, O., Liu, K., Rath, M., Ruebeta, D., Mueller, T., Skibbe, H., Drayer, B., Schmidt, T., Filippi, A., Nitschke, R., et al. (2012). ViBE-Z: a framework for 3D virtual colocalization analysis in zebrafish larval brains. Nat Methods 9, 735–742.

Rupp, B., Wullimann, M.F., and Reichert, H. (1996). The zebrafish brain: a neuroanatomical comparison with the goldfish. Anat Embryol (Berl) 194, 187–203.

Sajovic, P., and Levinthal, C. (1982). Visual cells of zebrafish optic tectum: mapping with small spots. Neuroscience 7, 2407–2426.

Schmitt., E.A., and Dowling., J.E. (1999). Early retinal development in the zebrafish, Danio rerio: light and electron microscopic analyses. J Comp Neurol 404(4), 515–536.

Schwartz, E.L. (1980). Computational anatomy and functional architecture of striate cortex: a spatial mapping approach to perceptual coding. Vision Res 20, 645–669.

Semmelhack, J.L., Donovan, J.C., Thiele, T.R., Kuehn, E., Laurell, E., and Baier, H. (2014). A dedicated visual pathway for prey detection in larval zebrafish. Elife 3.

Severi, K.E., Portugues, R., Marques, J.C., O’Malley, D.M., Orger, M.B., and Engert, F. (2014). Neural control and modulation of swimming speed in the larval zebrafish. Neuron 83, 692–707.

Simpson, J.I. (1984). The accessory optic system. Annu Rev Neurosci 7, 13–41.

Spillmann, L. (2014). Receptive fields of visual neurons: the early years. Perception 43, 1145–1176.

Tay, T.L., Ronneberger, O., Ryu, S., Nitschke, R., and Driever, W. (2011). Comprehensive catecholaminergic projectome analysis reveals single-neuron integration of zebrafish ascending and descending dopaminergic systems. Nat Commun 2, 171.

Thiele, T.R., Donovan, J.C., and Baier, H. (2014). Descending control of swim posture by a midbrain nucleus in zebrafish. Neuron 83, 679–691.

Trowe, T., Klostermann, S., Baier, H., Granato, M., Crawford, A.D., Grunewald, B., Hoffmann, H., Karlstrom, R.O., Meyer, S.U., Muller, B., et al. (1996). Mutations disrupting the ordering and topographic mapping of axons in the retinotectal projection of the zebrafish, Danio rerio. Development 123, 439–450.

Walley, R.E. (1967). Receptive fields in the accessory optic system of the rabbit. Exp Neurol 17, 27–43.

Wang, K., Hinz, J., Haikala, V., Reiff, D.F., and Arrenberg, A.B. (2019). Selective processing of all rotational and translational optic flow directions in the zebrafish pretectum and tectum. BMC Biol 17, 29.

Yamamoto, K., and Vernier, P. (2011). The evolution of dopamine systems in chordates. Front Neuroanat 5, 21.

Yanez, J., Suarez, T., Quelle, A., Folgueira, M., and Anadon, R. (2018). Neural connections of the pretectum in zebrafish (Danio rerio). J Comp Neurol.

Zhang, M., Liu, Y., Wang, S.Z., Zhong, W., Liu, B.H., and Tao, H.W. (2011). Functional elimination of excitatory feedforward inputs underlies developmental refinement of visual receptive fields in zebrafish. J Neurosci 31, 5460–5469.

Zimmermann, M.J.Y., Nevala, N.E., Yoshimatsu, T., Osorio, D., Nilsson, D.E., Berens, P., and Baden, T. (2018). Zebrafish Differentially Process Color across Visual Space to Match Natural Scenes. Curr Biol 28, 2018–2032 e2015.

